# A cell cycle-linked mechanism for the glutamine driven establishment of stem cell fate

**DOI:** 10.1101/2022.03.24.485713

**Authors:** Michael Xiao, Chia-Hua Wu, Graham Meek, Brian Kelly, Lyndsay E.A. Young, Sara Martire, Dara Buendia Castillo, Purbita Saha, Altair L. Dube, Matthew S. Gentry, Laura A. Banaszynski, Ramon C. Sun, Chintan K. Kikani

**Affiliations:** Department of Biology, College of Arts and Sciences, University of Kentucky, Thomas Hunt Morgan Building, 675 Rose Street, Lexington, KY 40506, USA; Molecular and Cellular Biochemistry, College of Medicine, University of Kentucky, BBS177, Lexington, KY 40536, USA; Cecil H. and Ida Green Center for Reproductive Biology Sciences, Children’s Medical Center Research Institute, Department of Obstetrics & Gynecology, Hamon Center for Regenerative Science and Medicine at University of Texas Southwestern Medical Center, Dallas, TX 75390, USA; Department of Neuroscience, College of Medicine, Department of Neuroscience, University of Kentucky, BBSRB Room B179, Lexington, KY 40536, USA; Weill Cornell/Rockefeller/Sloan Kettering Tri-Institutional MD-PhD Program, New York, NY 10021, USA

## Abstract

The cell cycle offers a unique opportunity for stem cells to sample metabolic and signaling cues to establish cell fate. Molecular pathways that integrate and convey these signals to cell cycle machinery to license cell fate transitions and drive terminal differentiation remain unknown. Here, we describe a signaling role of mitochondrial glutamine metabolism in driving exit from cell cycle-linked self-renewal to generate differentiation competent progenitors. In proliferating stem cells, mitochondrial glutamine metabolism opposes the WDR5-linked self-renewal network via acetylation and nuclear translocation of its upstream regulator, PASK. Nuclear PASK disrupts the mitotic WDR5-anaphase-promoting complex (APC/C) interaction to drive exit from self-renewal. Consistent with these roles, loss of PASK or inhibition of glutamine metabolism preserves stemness *in vitro* and *in vivo* during muscle regeneration. Our results suggest a mechanism whereby the proliferative functions of glutamine metabolism are co-opted by stem cells to establish cell fate.

## Introduction

Self-renewal is a property of stem cells where proliferation by cell division is coupled with the preservation of lineage identity and differentiation capabilities (stemness) (He et al., 2009). To preserve lineage identity, self-renewing stem cells must rapidly reactivate the expression of genes linked with cell identity following cell division as transcription is globally repressed during mitosis (Liu et al., 2017; Oh et al., 2020; Pelham-Webb et al., 2021). During terminal differentiation, self-renewing stem cells must exit the cell cycle and progressively generate a progenitor cell population that differentiates into functional cell types (Liu et al., 2019; Pauklin and Vallier, 2013). As self-renewing stem cells intrinsically preserve proliferation and oppose differentiation programs, metabolic and nutrient signals play critical roles in driving the transition from self-renewal to differentiation (Arnold et al., 2022; Baksh et al., 2020; Chakrabarty and Chandel, 2021; Chakraborty et al., 2019; Lu et al., 2021; TeSlaa et al., 2016).

Mitochondrial uptake of glutamine is thought to play an essential role in sustaining stem cell proliferation by generating ATP and maintaining redox balance (Yu et al., 2019). During muscle regeneration, glutamine secreted by macrophages sustains myoblast proliferation and promotes differentiation (Shang et al., 2020). However, whether metabolic signaling targets cell cycle-linked self-renewal processes to drive cell fate transitions remains unknown. Since the proliferative burst driven by glutamine metabolism is necessary for the expansion of the progenitor population (Yu *et al*., 2019), we set out to ask how metabolic pathways simultaneously couple proliferative processes with differentiation competence.

Using *in vitro* and *in vivo* myogenic differentiation as a model system, we show that glutamine metabolism drives heterogeneity in stem cell identity, as determined by Pax7 expression, facilitating the generation of differentiation competent progenitors. These functions of glutamine are driven primarily via acetylation and nuclear translocation of a stem cell enriched protein kinase, PASK, (Karakkat et al., 2019; Kikani et al., 2019; Kikani et al., 2016), which orchestrates biochemical disruption of mitotic self-renewal machinery required for the preservation of lineage identity. As glutamine is enriched early within the regenerating niche (Shang *et al*., 2020), which coincides with the peak of PASK expression and activity (Kikani *et al*., 2019; Kikani *et al*., 2016), we propose a signaling connection between glutamine metabolism and the signaling functions of PASK during tissue regeneration that results in exit from self-renewal and the generation of a progenitor population primed for differentiation.

## Results

### PASK inhibition preserves the stemness of mouse embryonic and adult stem cells

We have previously observed that PASK expression is positively correlated with proliferating (self-renewing) induced pluripotent, embryonic, and adult stem cells. It is rapidly downregulated following activation of the differentiation program in all systems (Kikani *et al*., 2016). Thus, as a metabolic sensory kinase enriched in stem cells (Karakkat *et al*., 2019; Kikani *et al*., 2019; Kikani *et al*., 2016), the study of PASK offers an opportunity to understand how cell fates are established in response to changing metabolic states. Functionally, PASK is required for the onset of terminal differentiation in mouse embryonic stem cells and adult MuSCs *in vitro* and *in vivo* (Kikani *et al*., 2016). However, its role and regulation in cycling stem cells undergoing self-renewal remains unknown. To understand the functional role of PASK in cycling stem cells, we cultured mouse embryonic stem cells (mESCs) in the 2i + LIF (2i) condition designed to maintain pluripotency and subsequently replaced the 2i media with PASKi (PASK inhibitor, BioE-1197, (Kikani *et al*., 2016; Wu et al., 2014)) + LIF (PASKi) to assess if PASKi can maintain pluripotency and stemness after the withdrawal of 2i. Strikingly, replacing 2i media with PASKi in mESCs resulted in a further increase in expression of genes associated with self-renewal and stemness (*Oct4, Sox2, and Rex1* mRNA) when compared with mESCs cultured in the 2iL conditions (Figure 1A, S1A). Additionally, using a Rex1-GFP reporter mESC line (Wray et al., 2011), we observed that cells cultured in PASKiL maintained GFP reporter expression at levels comparable to those observed from cells cultured in 2iL (Figure 1B, S1B). To determine how PASKi compares with 2i treatment in preserving the differentiation potential of mESCs, we performed an embryoid body (EB) formation assay of cells grown in 2iL or PASKiL. Cells grown in the 2iL culture condition differentiate well, as seen from the emergence of several fluid-filled cavitated structures after withdrawal of 2i. Strikingly, PASKiL cultured cells showed a substantial increase in the numbers and size of fluid-filled cavitated structures compared with 2i pretreated cells (∼52% for PASKi versus ∼12% for 2i treated cells, Figure S1C) after PASKi withdrawal. Consistent with our previous study, the presence of PASKi during EB formation attenuated differentiation as assessed morphologically (Figure S1C). Thus, our results indicate that PASK activity represents a novel target that modulates the decision between self-renewal and differentiation.

**Figure 1.**
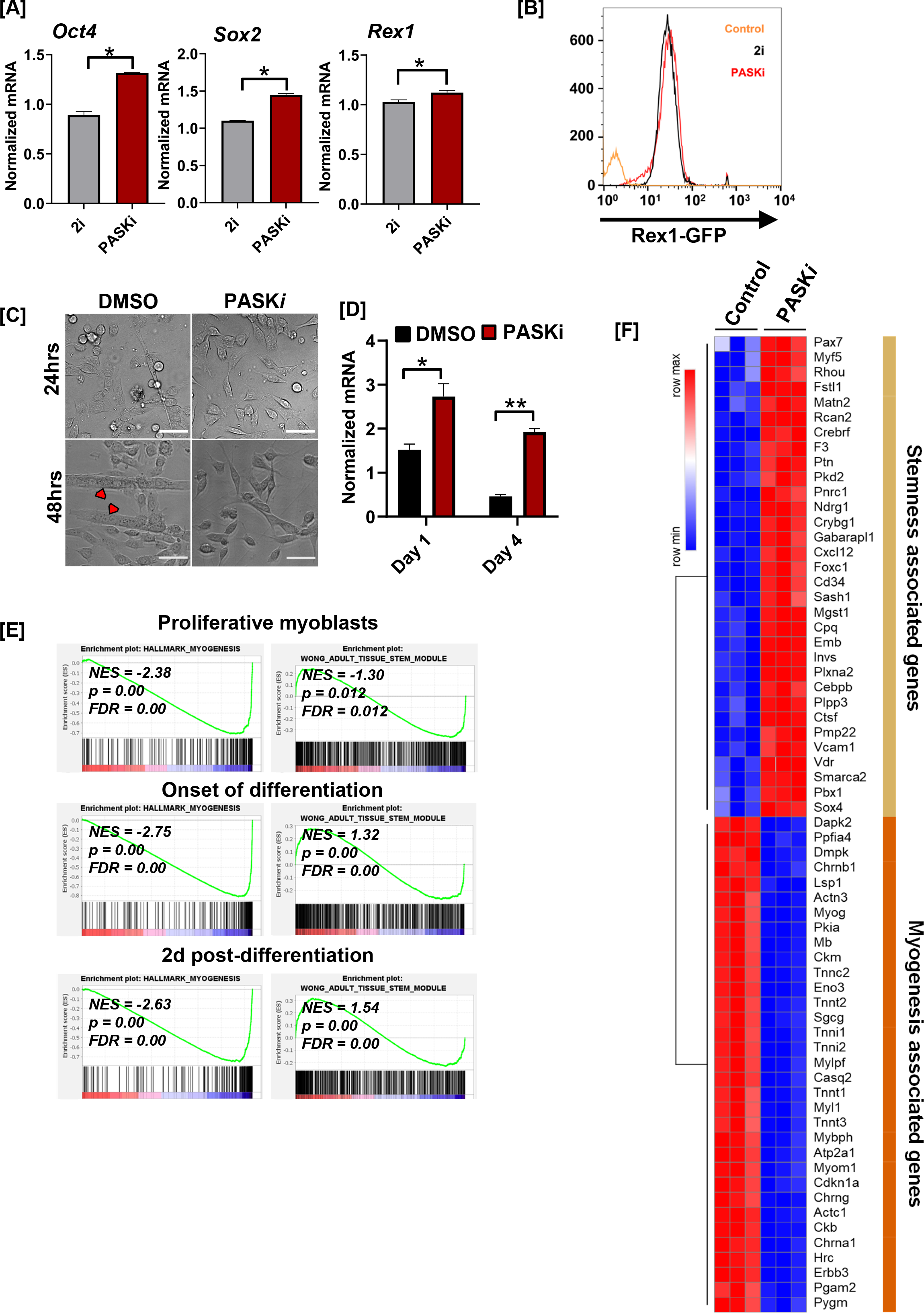
Loss of PASK function preserves stem cell identity. (A) RT-qPCR analysis of indicated transcript levels from 2i + LIF cultured mESCs after transitioning into 2i + LIF or PASKi + LIF conditions and cultured for 4 days. (B) Rex1-GFP intensity levels from mESCs cultured in 2i, vs. in PASKi as in (A). Rex1-GFP reporter expression was quantified by flow cytometry against a non-fluorescent control (Control). (C) Primary myoblasts were treated with DMSO or 50 µM PASKi for 4 days during the normal growth phase. Microscopy images were taken 24 hours or 48 hours post-isolation and treatment. Scale Bar = 40 µm. (D) Normalized (18s rRNA) mRNA levels of *Pax7* from primary myoblasts after 24 hours or 96 hours of culture in indicated conditions. (E) Gene set enrichment analysis of proliferating or differentiating C2C12 myoblasts by normalized enrichment scores for myogenesis and stemness. (F) RNA-sequencing analysis heatmap of differentially expressed genes associated with stemness and myogenesis in C2C12 myoblasts treated with DMSO (control) or PASKi for two days.

Since PASKi preserved mESC stemness, we asked if the same can be achieved for adult stem cells with PASKi treatment. PASK is expressed at very low levels in adult quiescent MuSCs. However, its expression increases rapidly as MuSCs are activated and begin to proliferate (Figure S1D)(Liu et al., 2013). To test how inhibition of mouse Pask during *in vitro* activation of MuSCs affects self-renewal and differentiation dynamics in adult stem cells, we cultured freshly isolated primary myoblasts in the presence or absence of PASKi. Pask inhibition was well-tolerated by isolated primary myoblasts, and PASKi treatment did not affect the proliferation rate of primary myoblasts. During extended *in vitro* culture, control myoblasts began to differentiate into nascent myotubes by 2 days post isolation and showed elongated myofibers by Day 4. On the other hand, PASKi treatment prevented precocious differentiation of primary myoblasts and preserved them in the proliferative state for a longer duration in culture (Figure 1C). Similar to our observations in mESCs where PASKi treatment increased the expression of stemness and pluripotency factors (Figure 1A), PASKi treatment of primary myoblasts resulted in increased transcript levels of *Pax7*, which is associated with MuSC self-renewal and myogenic lineage identity (Figure 1D).

Since primary myoblasts exhibit a rapid loss of self-renewal and undergo precocious differentiation when isolated from the muscle niche, the increase in *Pax7* mRNA levels could be attributed to an inhibition of differentiation by PASKi. Thus, we turned to non-transformed, cultured myoblasts such as C2C12, which can be maintained in a proliferative state for an extended duration under appropriate culture conditions. Using this system, we asked if PASKi treatment during the proliferative phase affects the expression of genes associated with stemness and self-renewal in cultured myoblasts. We performed RNA-seq analysis of C2C12 myoblasts treated with PASKi during proliferative (Day 0), early differentiating (Day 1), and late differentiating (Day 2) conditions. Our global transcriptomic analysis revealed a strong enrichment of genes associated with stemness and self-renewal at all time points when PASK is inhibited, starting with the proliferative condition (Figure 1E). Furthermore, PASK inhibition preserved self-renewal and stemness despite culture conditions that normally stimulate differentiation (Figure 1F, S1E-F). These results led us to hypothesize that PASK may play a crucial role in licensing exit from self-renewal and stemness preservation programs by regulating mechanisms that maintain the expression of genes associated with stemness (*Pax7* in MuSCs) and pluripotency (*Oct4* in mESCs).

### PASK is translocated to the nucleus at the onset of differentiation in muscle stem cells

The earliest event during differentiation is an exit from the stemness preservation program and the establishment of a committed progenitor population. Our results show that PASK plays an essential role in both processes (Figure 1, (Kikani *et al*., 2016)). To mechanistically understand PASK function in stem cells during this relatively short but crucial temporal window of early differentiation, we analyzed PASK subcellular distribution in proliferating versus differentiating myoblasts. In proliferating C2C12 myoblasts (Day 0), we find that the majority of the Pask is localized in the cytosol and excluded from the nucleus (Figure 2A). However, upon induction of the differentiation program, a significant fraction of Pask was localized to the nucleus as seen by overlapping intensities of the Pask signal with the nuclear marker DAPI and by biochemical fractionation (Figure 2A-B, S2A-B). We next investigated the relationship between Pask nuclear localization and its known signaling target, MyoG, a marker of early myogenesis (Kikani et al., 2016). We found a strong positive association between the presence of nuclear Pask and MyoG expression in proliferating and early differentiating primary myoblasts (Figure 2C-D). In contrast, the extent of nuclear Pask was inversely associated with the level of Pax7 (Figure 2E-F). Thus, under proliferating conditions, Pax7^hi^ (Figure 2E, yellow arrow indicate cells with stronger nuclear levels of Pax7) cells are more frequently associated with lower nuclear PASK presence, and Pax7^L0^ (Figure 2E, white arrows indicate cells with relatively weaker nuclear levels of Pax7) or Pax7^ab^ (cells lacking nuclear Pax7) are more frequently seen in cells with noticeable nuclear Pask presence (Figure 2F). Furthermore, we observed an asymmetrical subcellular distribution of Pask in primary myoblasts, where cells that retained nuclear Pask after mitosis also lacked Pax7 expression (Figure 2G). These data collectively establish a strong negative association between Pask nuclear localization and stem cell identity (Pax7 expression) in proliferating myoblasts.

**Figure 2.**
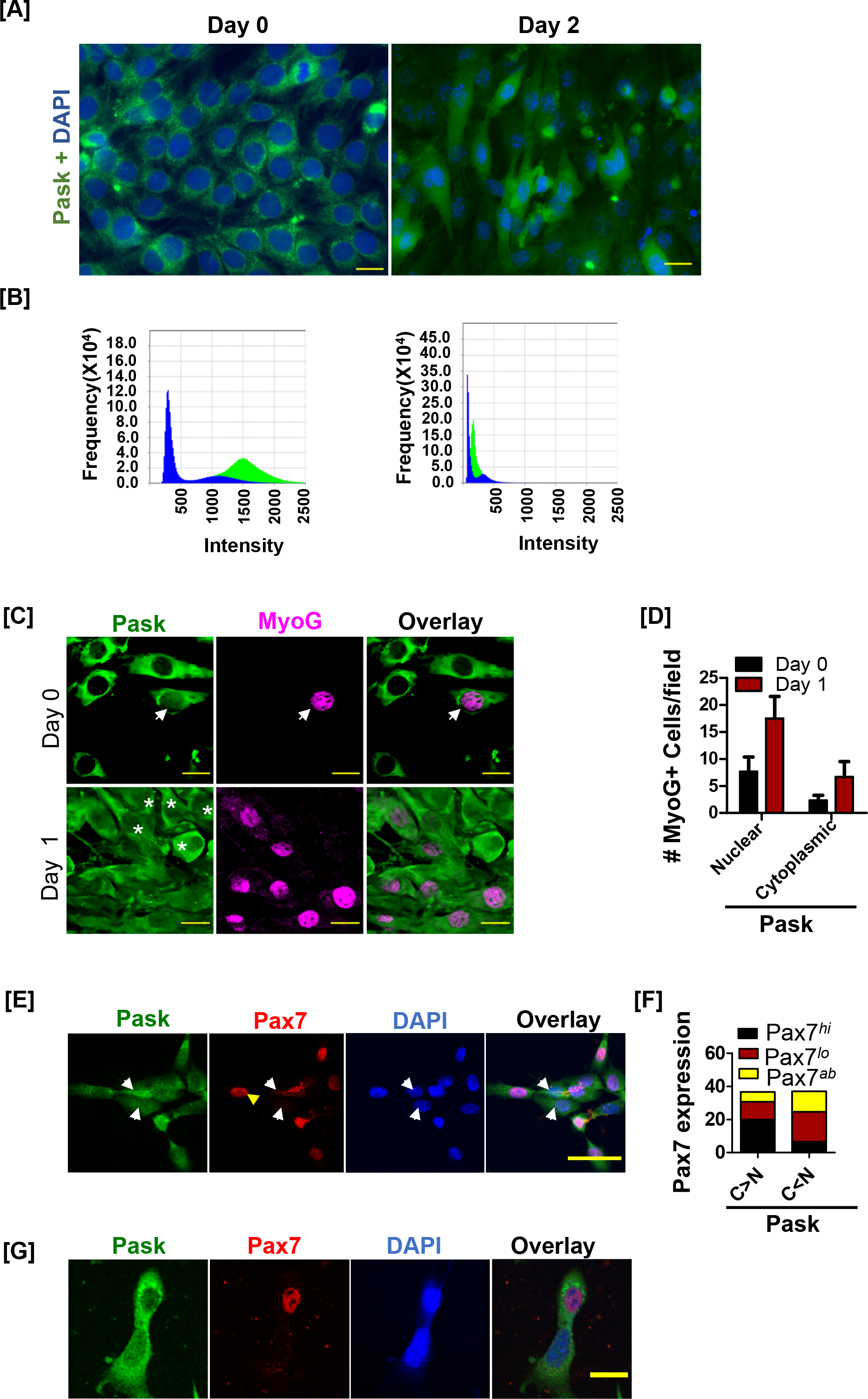
PASK is nuclear during differentiation and nuclear PASK downregulates stem cell identity. (A) Proliferating (day 0) or differentiating C2C12 (Day 2) myoblasts were stained with anti-Pask antibody. Scale Bar = 20 µm. (B) Immunofluorescence intensity overlays from the experiment in (A), showing overlap of Pask signal intensity (green) over nuclear signal (blue, DAPI). (C) Co-staining of Pask and MyoG from proliferating (day 0) and differentiating (day 1 post differentiation induction) primary myoblasts. Arrow indicates nuclear Pask expressing cells at Day 0. At Day 1, asterick corresponds to a subset of cells lacking nuclear Pask. Scale Bar = 20 µm. (D) The proportion of MyoG+ cells in nuclear vs. cytosolic Pask in proliferating (Day 0) or differentiating (Day 1) primary myoblasts. (E) Co-staining of Pask and Pax7 from proliferating (day 0) primary myoblasts. Arrow indicates strongly nuclear Pask expression in cells at Day 0. Scale Bar = 40 µm. (F) Distribution of Pax7 expression (high, low, or absent, as indicated by Pax7^hi^, Pax7^lo^, Pax7^ab^, respectively) in primary myoblasts showing more cytosolic (C>N) vs. more nuclear (C<N) Pask. (G) Asynchronously proliferating primary myoblasts were stained with Pask and Pax7 antibodies. Pax7 is excluded from daughter mitotic cells with asymmetric nuclear localization of Pask. Scale Bar = 40 µm.

### Signal regulated nuclear import-export machinery regulates nucleo-cytoplasmic shuttling of PASK

Because pharmacological inhibition of PASK stimulated Pax7 expression (Figure 1), we hypothesize that nuclear translocation of PASK is the cause and not the effect of loss of Pax7 expression. To test this hypothesis, we sought to first determine the mechanism of PASK translocation to the nucleus during early myogenesis. Previous high throughput studies have shown the interaction between human PASK and CRM1, the RanGTPase-driven exportin (Kırlı et al., 2015). Thus, we considered the possibility that PASK is a nucleo-cytoplasmic shuttling protein containing one or more nuclear export sequences (NES) and that regulated import and/or export may result in its nuclear localization at the onset of differentiation. To test this, we treated HEK293T cells expressing GFP-tagged WT human PASK (PASK) with a CRM1 inhibitor, Leptomycin B (LMB), under steady-state conditions. Surprisingly, LMB treatment did not result in a noticeable accumulation of PASK in the nucleus of HEK293T cells even at a high concentration (25nM) of LMB (Figure 3A). To determine if the nuclear import of PASK is rate-limiting in our LMB experiment, we fused a strong SV40-NLS (nuclear localization sequence) to GFP- PASK (NLS-PASK) to circumvent any mechanisms regulating nuclear import and retested its nuclear localization upon LMB treatment under the steady state. While the SV40-NLS fusion failed to drive PASK nuclear translocation on its own, LMB treatment induced a modest increase in nuclear localization of NLS-PASK (Figure 3A). Responsiveness of NLS-PASK to LMB treatment suggested the presence of one or more strong CRM1-dependent nuclear export sequences in PASK.

**Figure 3.**
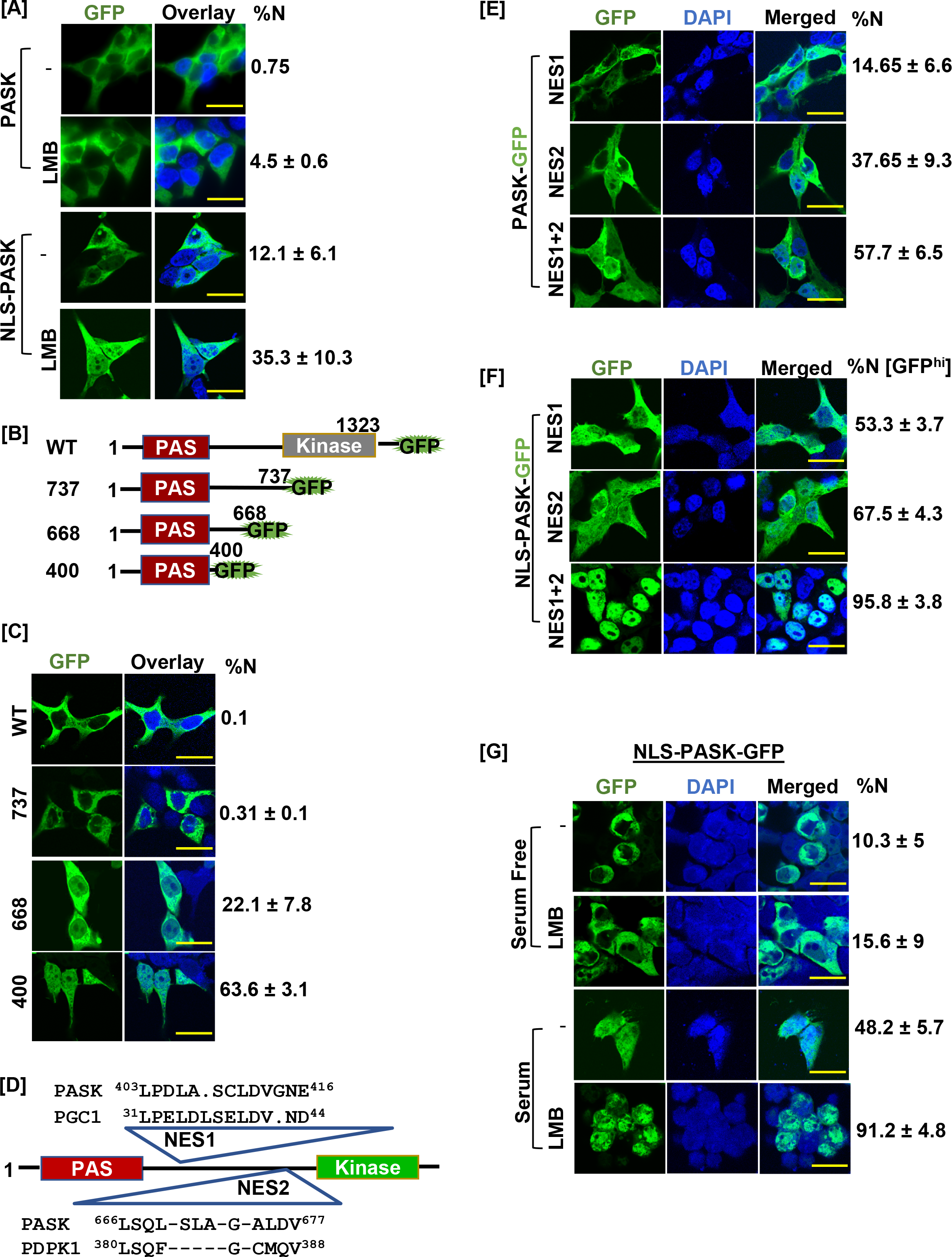
PASK nuclear translocation is governed by regulated nuclear export. (A) HEK-293T cells were transfected with GFP-tagged full-length human PASK (aa 1-1323, WT) or GFP-tagged full-length SV40 NLS-PASK (aa 1-1323, NLS-WT). Cells were treated with 25 nM Leptomycin B (LMB) for 2 hours as indicated, and analyzed by confocal microscopy. The percentage of cells containing any GFP signal in the nucleus in each condition was quantified (% cells showing nuclear (N±SD) GFP). All scale bars = 40 µm. (B) Domain illustration depicting GFP-tagged WT hPASK (aa 1-1323) and its truncated versions, GFP-tagged fragment 737 (aa 1-737), 668 (aa 1-668), and 400 (aa 1-400). PAS domain is highlighted in red. (C) HEK-293T cells were transfected with GFP-tagged full-length PASK (aa 1-1323, WT) and various truncations. Cells were analyzed by confocal microscopy. The percentage of cells containing any GFP signal in the nucleus was included in the quantification (%N±SD). (D) Diagram showing the locations of two nuclear export sequences (NES1 and NES2) and their sequence homology relative to PGC1 (NES1) and PDPK1 (NES2), respectively. (E) HEK-293T cells were transfected with GFP-tagged full-length WT PASK (aa 1-1323, WT) or NES1 (L^403^AL^405^S) or NES2 (L^666^SL^671^A). Cells were analyzed by confocal microscopy. The percentage of cells containing any GFP signal in the nucleus in each condition was quantified (% N±SD). (F) HEK-293T cells were transfected with GFP-tagged SV40 NLS-PASK (NLS-hPASK-GFP) containing mutated nuclear export sequences (NES1, NES2, or combined NES 1+2). Cells were analyzed by microscopy. The percentage of cells containing strong GFP signal (GFP^hi^) in the nucleus in each condition was quantified (% N±SD). (G) HEK-293T cells were transfected with GFP-tagged full-length SV40 NLS-hPASK (aa 1-1323, NLS-PASK-GFP). Cells were serum-starved (0.1 % serum) for 12 hrs and then subsequently stimulated with either 0.1 % serum (Serum Free) or 20 % serum (Serum) for 2 hrs. Cells were treated with 25 nM leptomycin B (LMB) where indicated and analyzed by confocal microscopy. The percentage of cells containing any GFP signal in the nucleus in each condition was quantified (% N±SD).

Proteins smaller than 60kDa could migrate to the nucleus by diffusion (Nigg, 1997). With this size limitation in mind, we performed a series of C-terminal truncations in PASK to identify the region that mediates PASK nuclear export (Figure 3B). Scoring for cells showing at least some nuclear GFP presence, we found that the GFP-1-737 (MW = ∼115kDa) fragment remained predominantly cytoplasmic (Figure 3C). A smaller fragment, GFP-1-660 (MW = ∼100kDa) showed nuclear GFP expression in ∼22 % of cells (Figure 3C), while fragment GFP-1-400 (MW = ∼ 70kDa) showed increased nuclear localization of PASK with ∼67 % of cells showing at least some nuclear GFP presence (Figure 3C). These results suggested the presence of nuclear export sequences between amino acids 400 and 660 and between amino acids 660-737. We used multiple bioinformatic tools to identify L401-L409 (NES1) and L666-L671 (NES2) as putative NES sequences in PASK (Figure S3A-C, Figure 3D) (Xu et al., 2015; Xu et al., 2021). The NES1 residues are similar to an experimentally verified NES in PGC1-α (Chang et al., 2010) and the NES2 residues are similar to the nuclear export sequence of PDK1 that we previously discovered (Figure 3D) (Lim et al., 2003). We found that mutation of either NES1 or NES2 residues resulted in a modest but statistically significant increase in the number of cells with nuclear PASK localization compared to WT-PASK or NLS-PASK in the absence of LMB (Figure 3E). Combined mutation of NES1 and NES2 resulted in a significantly increased proportion of cells that contain nuclear PASK, with the extent of nuclear PASK in each cell similar to what we observed for PASK during myogenesis (Figure 3E). However, incomplete nuclear retention of NES1+NES2 mutated PASK prompted us to examine if the nuclear import of PASK might be rate-limiting, preventing stronger PASK nuclear accumulation despite NES1 and NES2 mutations. To test this, we generated separate NES1 or NES2 or NES1+NES2 combined mutations in NLS-PASK. While mutations of NES1 or NES2 in NLS-PASK improved nuclear retention of NLS-PASK (Figure 3F), mutating both NES1 + NES2 in NLS-PASK resulted in robust nuclear retention of NLS-PASK in nearly 100 % of cells (Figure 3F). These results conclusively show that NES1 and NES2 are functional for exporting PASK from the nucleus.

Since the nuclear accumulation of PASK was induced by differentiation signaling cues (Figure 2), we asked if signaling pathways target the nuclear export machinery (NES sequences) to retain PASK in the nucleus. To test this, we used NLS-PASK to circumvent any regulatory import mechanisms that may be present. We serum-starved HEK293T cells expressing NLS-PASK and treated them with LMB in the presence or absence of acute serum stimulation for 2 hrs. Remarkably, NLS-PASK was completely excluded from the nucleus in serum-starved cells despite LMB treatment (Figure 3G). In contrast, acute serum stimulation resulted in increased nuclear accumulation of NLS-PASK even without LMB (Figure 3G). Furthermore, serum stimulation improved the effectiveness of LMB treatment, as seen from stronger nuclear retention of PASK (in nearly 100 % of cells), similar to what was observed with NES1+NES2 mutated NLS-PASK. The improved effectiveness of LMB in blocking PASK nuclear export suggests that serum stimulation weakens the effects of the NES1 and/or NES2 sequences to reduce the nuclear export rate of PASK.

### Glutamine-dependent acetylation of PASK stimulates its nuclear accumulation

We next asked if WT-PASK lacking SV40-NLS could also be translocated to the nucleus in response to serum stimulation. As shown in Figure 4A, we observed significant nuclear accumulation of GFP-PASK by 2 hrs of serum stimulation. In addition, endogenous PASK was also translocated to the nucleus in response to acute serum stimulation in C2C12 myoblasts (Figure S4A). We have previously shown that PASK is phosphorylated and activated by mTORC1 in response to serum stimulation (Kikani *et al*., 2019). This phosphorylation activates PASK to stimulate the myogenesis program. Two of the mTOR phosphorylation sites on PASK, Ser^640,^ and Thr^642^, are situated near NES2 (L666-L671) residues. Therefore, we tested if mTOR signaling mediates the nuclear translocation of WT-PASK in response to serum stimulation. However, inhibition of mTORC1 by rapamycin did not consistently or significantly block serum-induced nuclear translocation of WT-PASK, suggesting that mTORC1 signaling alone is not sufficient for driving PASK nuclear translocation in response to serum stimulation (Figure 4A).

**Figure 4.**
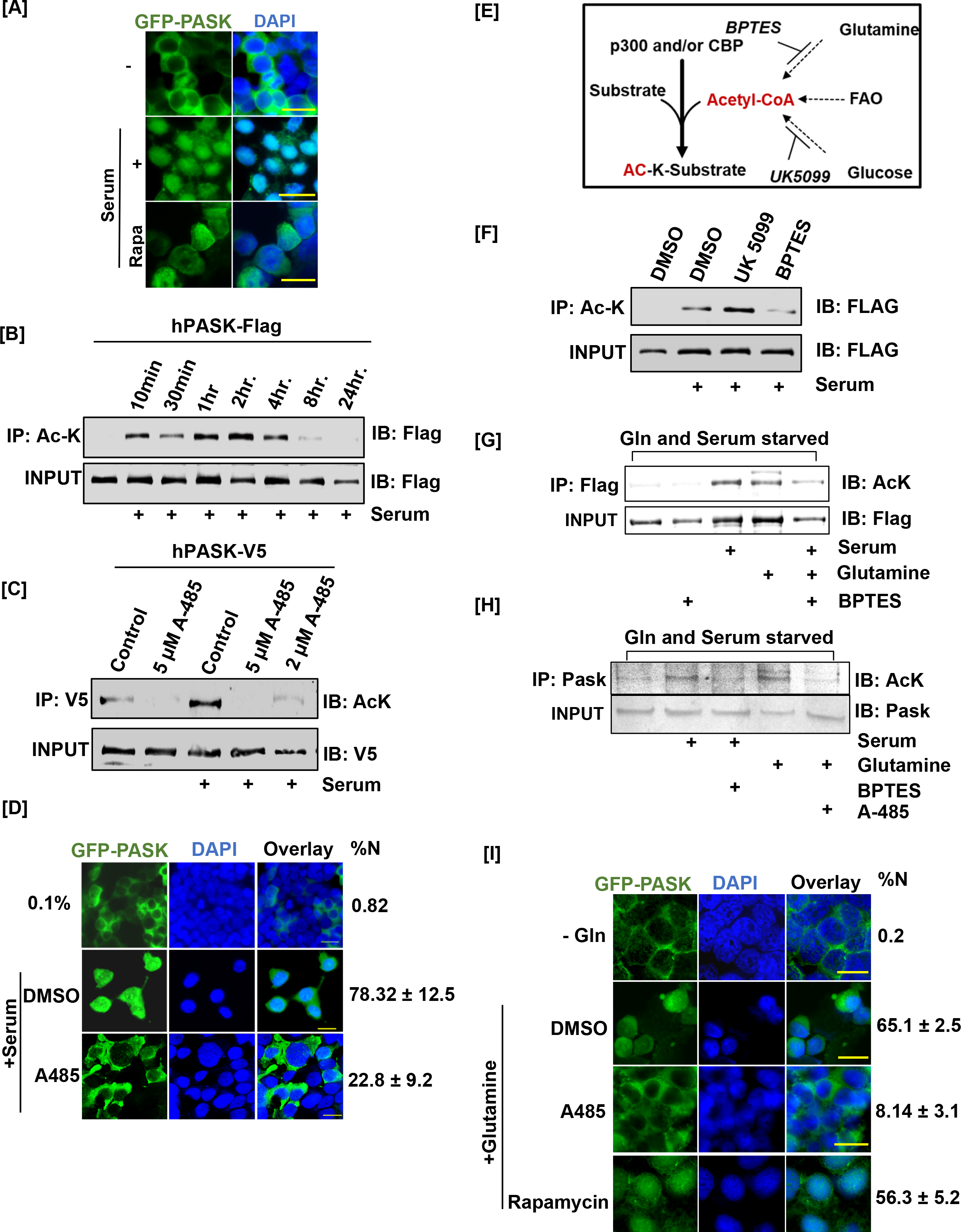
Glutamine driven acetylation promotes PASK nuclear translocation. (A) HEK-293T cells stably expressing GFP-tagged full-length WT PASK (aa 1-1323) were serum- starved (0.1% serum) for 12 hours and then stimulated for 2 hours with 20% serum. For the rapamycin- treated condition, cells were pretreated with 100 nM rapamycin for 2 hrs in serum-free conditions prior to 20 % serum induction. Scale Bar = 40 µm. (B) HEK-293T cells stably expressing FLAG-tagged full-length PASK (aa 1-1323) were serum- starved (0.1 % serum) for 12 hours. Cells were subsequently stimulated with either 0.1 % serum or 20 % serum for the time intervals indicated. Total cellular acetylome was purified from cells using anti-Ac-K antibody and the presence of acetylated PASK was detected by anti-Flag antibody. (C) V5-tagged full-length PASK (aa 1-1323) was transiently expressed in HEK-293T cells. 24 hrs after transfections, cells were serum-starved in 0.1% serum for 12 hours. Cells were pretreated with 5 μM or 2 μM A-485 to inhibit p300/CBP acetyltransferases one hour prior to serum stimulation. Cells were stimulated with either 0.1 % serum or 20 % serum for 2 hours. V5-tagged PASK was immunoprecipitated from cells, and the presence of acetylated PASK was detected using an anti-Ac-K antibody. (D) HEK-293T cells stably expressing GFP-tagged full-length PASK (aa 1-1323) were serum-starved (0.1 % serum) for 12 hours. Cells were pretreated with 2 μM A-485 to inhibit p300/CBP acetyltransferases one hour prior to serum stimulation. Cells were stimulated with either 0.1 % serum (-) or 20 % serum (Serum) for 2 hours, and the extent of nuclear GFP-tagged PASK (% N±SD) was analyzed by immunofluorescence. Scale Bar = 20 µm. (E) Illustration of metabolic pathways involved in acetyl-CoA production that can fuel cellular acetylation. (F) HEK-293T cells stably expressing FLAG-tagged full-length PASK (aa 1-1323) were treated with 10 µM UK 5099 (mitochondrial pyruvate carrier inhibitor) or 10 µM BPTES (glutaminolysis inhibitor) overnight along with 0.1% serum starvation where indicated. Cells were stimulated with 20 % serum in the presence of DMSO or indicated inhibitors. hPASK was purified using anti-Flag antibody and probed with anti-AcK antibody. (G) C2C12 cells stably expressing FLAG-tagged full-length PASK (aa 1-1323) were serum starved in glutamine-free media for 12 hours. 10 µM BPTES was added along with serum/glutamine starvation where indicated and maintained during 2 hrs serum stimulation. For glutamine stimulation, media containing 2 mM glutamine was added with or without 10 µM BPTES. hPASK was purified using anti- Flag antibody and probed with anti-AcK antibody. (H) Primary myoblasts were serum and glutamine starved for 12 hrs in the presence of 10 µM BPTES as indicated. Cells were stimulated with 20 % serum or 2 mM glutamine in the presence of either 2 μM A-485 (for glutamine stimulated cells) or 10 µM BPTES (for serum stimulated cells) for 2 hours. The lysate was immunoprecipitated for endogenous PASK prior to immunoblotting for total acetyl-lysine (Ac- K). (I) HEK-293T cells stably expressing GFP-tagged full-length PASK (aa 1-1323) were serum and glutamine starved for 12 hours. Cells were pre-treated with DMSO (as control), 100 nM rapamycin or 2 μM A-485 for 2 hrs prior to stimulation with 2 mM glutamine. Cells were fixed and the extent of nuclear GFP-tagged PASK was quantified by confocal microscopy (%N±SD). Scale Bar = 40 µm.

Along with phosphorylation, acetylation of non-histone proteins plays important roles in controlling nucleo-cytoplasmic shuttling and cell fate decisions in stem cells (Choudhary et al., 2014). To determine if endogenous PASK is acetylated, we treated C2C12 myoblasts with Trichostatin A (TSA), a broad-spectrum histone deacetylase inhibitor. Probing the purified acetylome with an anti-PASK antibody revealed strong enrichment of acetylated endogenous PASK (Figure S4B). We also noticed a dose (Figure S4C) and time-dependent (Figure 4B) increase in PASK acetylation caused by serum stimulation in HEK293T cells.

To examine the physiological relevance of PASK acetylation *in vivo,* we injured tibialis anterior (TA) muscles of *Pask*^het^ mice and analyzed PASK acetylation at day 3 and day 5 post-injury. We noticed that a significant fraction of mouse Pask is acetylated during regeneration at days 3 and 5 post-injury (Figure S4D). Collectively, these results show that Pask is a novel non-histone protein target of signal-dependent acetylation in cells and tissues.

Due to the overlapping temporal kinetics of PASK acetylation (Figure 4B) and nuclear translocation (Figure 3G, 4A), we asked whether PASK acetylation drives its nuclear translocation. For this, we first sought to identify the upstream histone acetyltransferase (HAT) that catalyzes PASK acetylation in response to serum stimulation. KAT2A and p300/CBP are two major histone acetyltransferases upregulated in MuSCs during regeneration and are known to induce acetylation of non-histone proteins during myogenesis (Das et al., 2017; Puri et al., 1997a; Puri et al., 1997b; Sartorelli et al., 1999). Whereas siRNA-mediated knockdown of KAT2A did not affect PASK acetylation induced by serum (Figure S4E), pretreatment of cells with a highly selective p300/CBP inhibitor, A-485 (Lasko et al., 2017), nearly completely blocked serum-induced PASK acetylation (Figure 4C) and serum-induced PASK nuclear localization (Figure 4D). Thus, p300/CBP mediated acetylation of PASK stimulates its nuclear translocation.

Serum-induced PASK acetylation was independent of inputs from signaling kinases, as inhibition of PI-3K, mTOR, or Akt did not prevent PASK acetylation induced by serum (Figure S4F). Therefore, we explored the involvement of metabolic pathways in driving PASK acetylation. Acetyl-CoA is generated via glucose (via mitochondrial pyruvate oxidation) and glutamine (via glutaminolysis). Strikingly, inhibition of the mitochondrial pyruvate complex (MPC1) by UK5099 resulted in more robust induction of PASK acetylation than serum alone, perhaps due to a compensatory increase in glutaminolysis (Figure 4F) (Yang et al., 2014). Consistent with this, inhibition of mitochondrial glutaminolysis by a GLS1 specific inhibitor, BPTES, resulted in a near-complete loss of serum-induced PASK acetylation (Figure 4F). Furthermore, glutamine addition alone was sufficient to stimulate PASK acetylation as much as serum induction in serum and glutamine starved conditions (Figure 4G). The addition of BPTES significantly reduced PASK acetylation induced by combined serum and glutamine stimulation (Figure 4G). Finally, in isolated primary myoblasts, glutamine metabolism drives acetylation of the endogenous Pask via p300/CBP since glutamine-stimulated Pask acetylation was blocked by pretreatment with the p300/CBP inhibitor, A-485 (Figure 4H).

We next asked if glutamine stimulation is sufficient to cause PASK nuclear localization. As shown in Figure 4I, glutamine starvation resulted in the nuclear exclusion of PASK. The addition of glutamine stimulated robust nuclear translocation of PASK, which was blocked by p300/CBP inhibition (A-485), but not by mTORC1 inhibitor, rapamycin (Figure 4I). Thus, our results show that glutamine signaling to p300/CBP, but not mTOR, regulates PASK nuclear translocation via acetylation.

### Selective glutamine withdrawal triggers the preservation of stemness in muscle stem cells

The critical role of glutamine in stimulating PASK nuclear translocation prompted us to examine the role of glutamine metabolism in various stages of self-renewal and differentiation. As expected, based on previous reports (Ahsan et al., 2020; Shang *et al*., 2020), glutamine withdrawal in cultured myoblasts significantly reduced proliferation (Figure 5A). The glutamine withdrawn cells were significantly enlarged and remained alive in cell culture for at least 4 days (Figure 5A). Surprisingly, acute glutamine withdrawal also resulted in a robust increase in the proportion of Pax7+ myoblast numbers (Figure 5B-C).

**Figure 5.**
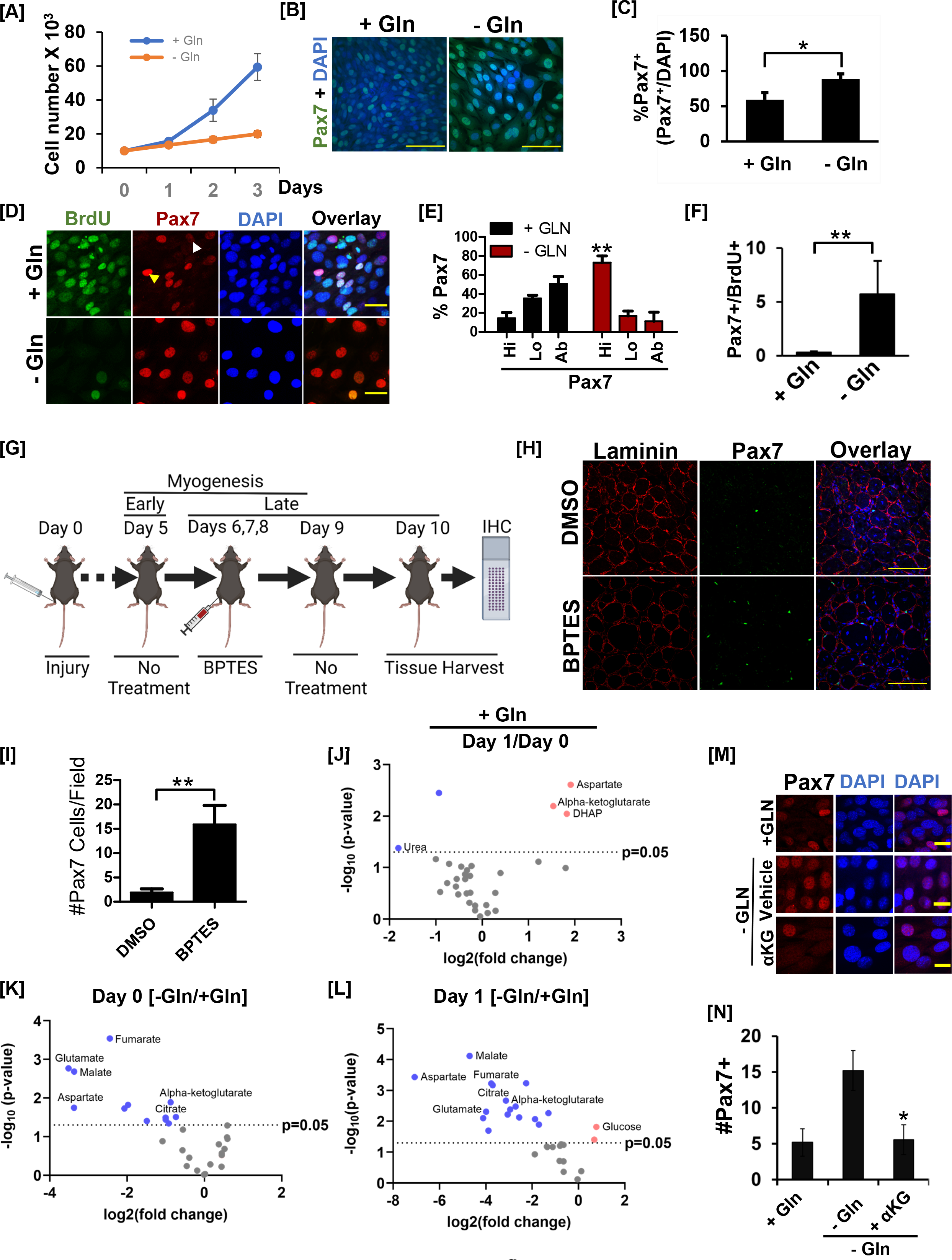
Inhibition of glutamine metabolism preserves stem cell identity. (A) Cell proliferation rate of C2C12 myoblasts cultured in the presence (+ Gln) or absence (- Gln) of glutamine over 3 days. 10^4^ cells were seeded in 60 mm tissue culture dishes. 24 hrs after seeding, cells were transferred to media containing 10 % serum with or without 2 mM Glutamine. Cells were counted every day for 3 days using an automated cell counter. (B) C2C12 myoblasts were allowed to proliferate in the presence (+ Gln) or absence (- Gln) of glutamine. Relative levels of Pax7 were visualized by immunofluorescence microscopy. Scale Bar = 100 µm. (C) Quantification of Pax7+ cell numbers in the presence (+ Gln) or absence of glutamine (- Gln). (D) 48 hours after isolation and culture in media containing glutamine, primary myoblasts were allowed to proliferate in the presence (+ Gln) or absence (- Gln) of glutamine for 3 days. Cells were pulsed with 10 µM BrdU in + Gln or - Gln media for 24 hrs before cell fixation. Cells were fixed and BrdU incorporation and Pax7+ cell numbers were analyzed by immunofluorescence microscopy using antibodies against BrdU and Pax7. Scale Bar = 40 µm. (E) Numbers of cells showing Pax7 expression at high (hi), low (lo) or absent (ab) levels in the presence (+ Gln) or absence (- Gln) of glutamine. ** *P<0.005* (% Pax7^hi^ cells in - Gln vs. + Gln). (F) Quantification of immunofluorescence analysis from (D) for Pax7^+^/BrdU^+^ in isolated muscle stem cells cultured in the presence (+ Gln) or absence (- Gln) of glutamine. (G) Experimental set up to test the effect of acute Gls1 inhibition by BPTES injection on Pax7 levels. 12 weeks old C57BL/6 mice were injured with 25 µl of 1.2 % w/v BaCl2 to the tibialis anterior (TA) muscles. On days 6, 7, and 8 post-injury, 10 μl of PBS containing 0.01 % DMSO or 10 μM BPTES was injected intramuscularly into the belly of TA muscles. Tissues were harvested on day 10. Tissue sections were stained for Laminin (red) or Pax7 (green) and analyzed by confocal microscopy. (H) Immunofluorescence images of muscle sections stained with Pax7 and Laminin from experiment in (G). Scale Bar = 100µm. (I) Quantification of average Pax7+ cell number/field from images in (H). (J) Volcano plot of metabolite abundances in differentiating (day 1) versus proliferating (day 0) C2C12 myoblasts in the glutamine replete condition. (K) Volcano plot of metabolite abundances in proliferating (day 0) C2C12 myoblasts cultured in the presence (+ Gln) or absence (- Gln) of glutamine. (L) Volcano plot of metabolite abundances in differentiating (day 1) C2C12 myoblasts cultured in the presence (+ Gln) or absence (- Gln) of glutamine. (M) C2C12 myoblasts were grown in the presence (+ Gln) or absence of glutamine (-Gln) for 24 hrs. 1 mM cell-permeable dimethyl-alpha ketoglutarate (aKG) or vehicle control was added to glutamine depleted cells for 24 hrs. After treatment, cells were fixed and stained with anti-Pax7 antibody. (N) Quantification of Pax7+ cell number from experiment in (M). Scale Bar = 20 µm.

Since progression through the cell cycle is linked with the preservation of stem cell identity during self-renewal, our data suggest that glutamine depletion activates the stemness preservation program when the proliferation of stem cells is metabolically unfavorable. To test this, we determined the extent of Pax7 nuclear staining in BrdU+ and BrdU- populations of isolated primary myoblasts under normal or glutamine depleted culture conditions. BrdU+ cells (proliferating cells) in glutamine-rich media were smaller in size and displayed greater heterogeneity in the level of nuclear Pax7 expression (Figure 5D-E). We also noticed a significant Pax7^L0^ and Pax7^ab^ population among the BrdU+ cycling stem cells (Figure 5D-E). In contrast, glutamine withdrawal blunted BrdU incorporation yet resulted in a robust increase in the proportion of Pax7^hi^ cells and Pax7 staining intensity within individual cells (Figure 5D- E). Since the ratio of Pax7+/BrdU+ nuclei is increased by glutamine withdrawal (Figure 5F), our results indicate that glutamine withdrawal results in a reactivation of Pax7 expression in Pax7^lo^ muscle stem cells. Consistent with this, *Pax7* mRNA levels were significantly increased when glutamine was withdrawn (Figure S5A). In addition to *Pax7*, transcript levels for other stemness and self-renewal associated genes, such as *Cd34*, *Myf5*, and *Myod*, were also significantly elevated upon glutamine deprivation (Figure S5A). Furthermore, we also noticed a significant increase in the expression of *Foxo3*, a marker for MuSC quiescence (Figure S5A), while expression of the proliferation marker *Ki67* was nearly completely abolished in glutamine-depleted cells (Figure S5A). These results suggested that glutamine withdrawal enforces the stemness preservation program and induces quiescence *in vitro* in a cell-autonomous manner. Finally, glutamine withdrawal also resulted in a near-complete loss of *Myog*, indicating a dependence on glutamine metabolism to generate the committed progenitor population (Figure S5A). We validated these functions of glutamine in cultured myoblasts as glutamine withdrawal or inhibition of Gls1 by BPTES treatment increased Pax7 mRNA levels in C2C12 myoblasts (Figure S5B). Furthermore, while glutamine depletion blocked myogenesis, inhibition of mitochondrial pyruvate import by UK5099 strongly stimulated differentiation as measured by *Myog*, *Mylpf*, and *Acta1* transcript levels (Figure S5B) and immunostaining (Figure S5C), perhaps via increased glutaminolysis (Yang *et al*., 2014).

During the early muscle regeneration response, glutamine, enriched in the regenerating niche, is taken up by activated muscle stem cells which drives their proliferation and is required for myogenic commitment and the regenerative myogenesis program (Shang *et al*., 2020). According to a recent report, a quiescent, self-renewing population is established after this early wave of myogenesis program has largely concluded (Day 5-7 post-injury) (Cutler et al., 2021). Our results show that acute glutamine withdrawal resulted in reactivation of *Pax7* transcription, decrease in proliferation, and increased expression of the quiescence marker, *Foxo3*, in isolated MuSCs. Therefore, we asked if loss of glutamine signaling could contribute to the reestablishment of Pax7+ stem cells and the expansion of the self-renewing stem cell population after the early wave of regenerative myogenesis program is completed. To test this, we injured the tibialis anterior (TA) muscles from WT mice and allowed them to undergo regenerative myogenesis up to Day 5. This ensures normal progression through the early myogenesis program. On Day 5, we began to intramuscularly inject 10 µM BPTES or DMSO every day for three days, followed by one day of no additional treatment to allow for clonal expansion of cell populations.

We then harvested TA muscles and quantified Pax7+ stem cell numbers (Figure 5G). Treatment of BPTES after early myogenesis did not significantly affect regenerative myofiber sizes (Figure 5H). In contrast, we saw a significant increase in Pax7+ cell numbers in BPTES compared with DMSO-treated animals (Figure 5H-I). Taken together, our results suggest that proliferative functions glutamine metabolism are linked with stemness and differentiation decisions *in vitro* and *in vivo*.

We next sought to understand the metabolic function of glutamine in balancing stemness with cell proliferation. Compared with proliferating cells, early differentiating myoblasts show significantly increased intracellular glutamine levels and their associated downstream metabolites, glutamate, alpha-ketoglutarate, and aspartate (Figure 5J). We were surprised by this observation since early differentiating myoblasts have already exited the cell cycle, while increased glutamine metabolism is linked with proliferative processes. Similarly, under glutamine withdrawn conditions where proliferation is inhibited, but stemness is enhanced (Figure 5A-D), we noticed a significant reduction in alpha-ketoglutarate, citrate, and malate levels (Figure 5K-L, S5D-E). Thus, glutamine metabolism in adult stem cells could serve to couple proliferation with the cell fate transition to favor differentiation. Consistent with this hypothesis, we find that the addition of cell-permeable alpha-ketoglutarate reversed the increased Pax7+ cell number due to glutamine withdrawal (Figure 5M-N). Taken together, our results suggest that glutamine metabolism driven proliferation plays a key role in driving heterogeneity in stem cell identity (via Pax7 expression) to generate a differentiation-competent progenitor population.

### Glutamine disrupts the WDR5-APC/C interaction during mitosis to drive exit from self-renewal via PASK nuclear translocation

As a downstream target of glutamine metabolism, we asked if PASK mediates the effects of glutamine metabolism on Pax7 levels and cell identity. To test this, we treated proliferating primary myoblasts with PASK inhibitor (PASKi). Unlike glutamine withdrawal, PASK inhibition did not affect cell proliferation rate or size (Figure S6A). Strikingly, similar to glutamine withdrawal (Figure 5D-E), PASK inhibition significantly increased Pax7+ myoblast numbers and reduced heterogeneity of its nuclear staining intensities across the field (Figure 6A, B). We next asked if loss of PASK also affects Pax7+ stem cell number *in vivo* during muscle regeneration. Interestingly, *Pask*^-/-^ animals showed a marked increase in Pax7+ cell numbers as early as Day 5 post-injury, which remained significantly elevated even at Day 14 when Pax7 numbers in *Pask*^+/+^ animals were significantly reduced (Figure 6C- D). Thus, loss of PASK or withdrawal of glutamine signaling results in an increased proportion of Pax7+ myoblast numbers *in vitro* and Pax7+ MuSC numbers during muscle regeneration.

**Figure 6.**
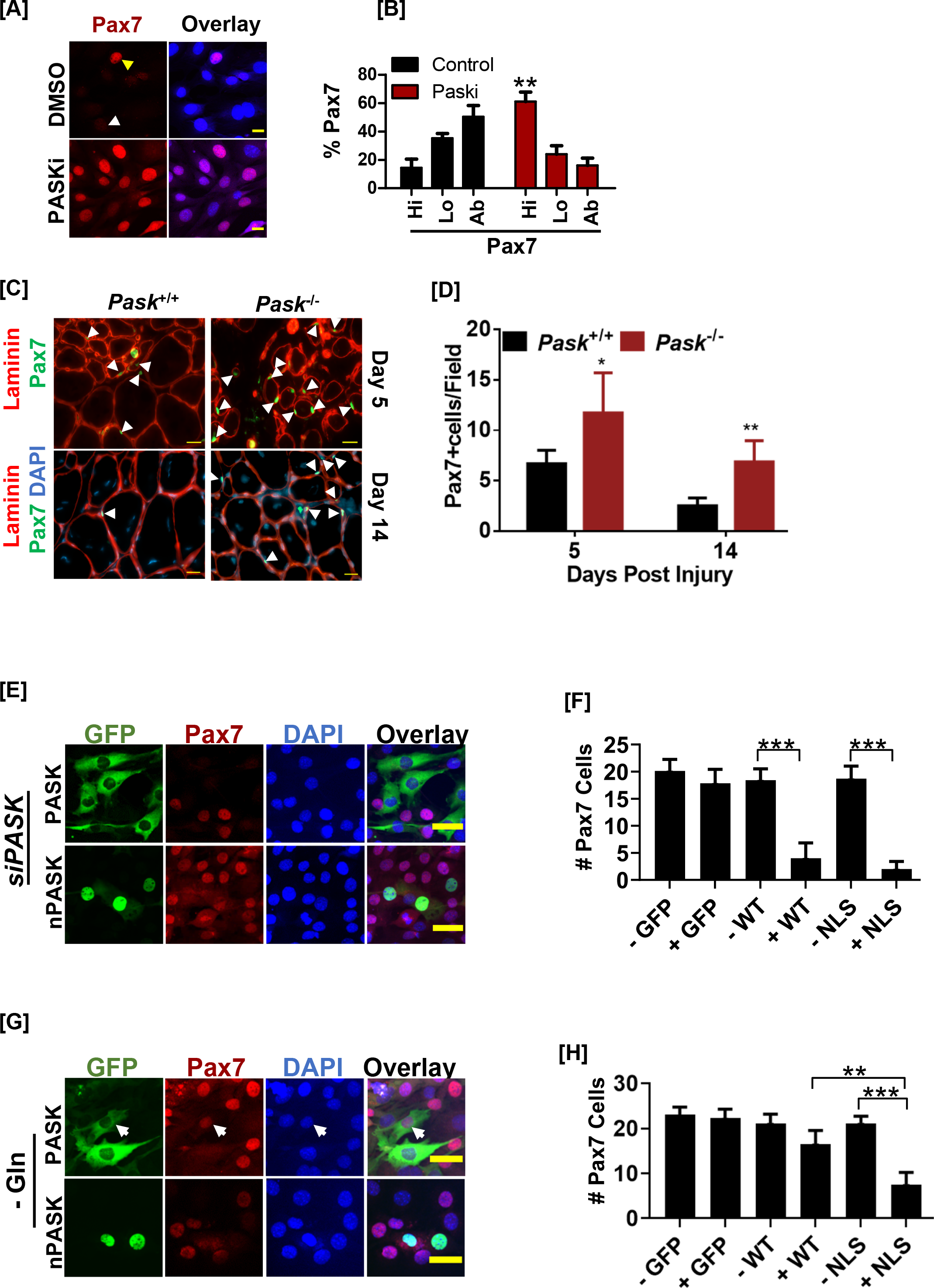
Glutamine driven establishment of cell identity requires nuclear PASK function. (A) Pax7+ cells were visualized by staining primary myoblasts after 2 days of treatment with DMSO or 50 µM Paski. Scale Bar = 20µm. (B) Numbers of cells showing Pax7 expression at high (hi), low (lo), or absent (ab) levels from DMSO or 50 µM Paski treated cells. (C) TA muscles of littermates of *Pask^WT^* and *Pask^KO^* animals were injured using 1.2 % BaCl2. Frozen tissue sections were stained using anti-Pax7 and anti-Laminin antibodies. Arrows indicate examples of Pax7+ cells identified and used for quantification in (D). Scale Bar = 40 µm. (D) Quantification of Pax7+ MuSCs numbers at Day 5 or Day 14 in *Pask^WT^* vs. *Pask^KO^* mice. Scale bars = 20 μm. n=3. (E) Mouse Pask was silenced using multiplex siRNAs in C2C12 cells. 24 hrs after siRNA transfection, GFP-tagged WT or NLS-hPASK^L403L405S+L666S-L671A^-hPASK (nuclear PASK, nPASK) was retrovirally expressed. 48 hrs after viral infection, cells were fixed and Pax7+ cells were analyzed by immunofluorescence against Pax7. Pask expressing cells were identified by GFP expression. Scale Bar = 40µm. (F) Quantification of Pax7^hi^ cell numbers in individual cells expressing GFP control (GFP+), WT- PASK (WT+) or nuclear PASK (nPASK+) compared with uninfected cells (GFP-, WT- or nPASK-) from experiment in (E). (G) Pax7 expression was assessed by immunofluorescence microscopy from C2C12 myoblasts cultured for 48 hours in glutamine-depleted conditions expressing GFP control, GFP tagged WT or nPASK plasmids. Scale Bar = 40 µm. (H) Quantification of Pax7^hi^ cell numbers in individual cells expressing GFP control (GFP+), WT- PASK (WT+) or nuclear PASK (nPASK+) compared with uninfected cells (GFP-, WT- or nPASK-) from experiment in (G).

Parallels between glutamine withdrawal or PASK inhibition prompted us to inquire if nuclear PASK is essential for glutamine metabolism to regulate cell identity. We first tested the ability of cytosolic (WT) versus nuclear PASK (nPASK; NLS^NES1+2^-PASK) in downregulating Pax7 expression in PASK silenced cells in the glutamine replete condition. Under this condition, Pax7+ cell numbers and the relative intensities of Pax7 staining increased in the cell population. Expressing either WT or nuclear PASK, but not GFP control, effectively reverses the number of Pax7^hi^ cells observed due to *Pask* silencing (Figure 6E-F, Figure S6B). However, under glutamine-depleted conditions, nuclear PASK was significantly more effective than WT-PASK or GFP control at reversing the number of Pax7^hi^ myoblasts (Figure 6G-H, Figure S6B).

Our results show that the expression of stemness-related genes increases with either glutamine depletion or PASK inactivation. Therefore, we asked whether glutamine and PASK act upon the same downstream targets. We and others have previously identified WDR5 as a PASK interacting protein and its substrate (Karakkat *et al*., 2019; Kikani *et al*., 2016). We showed that the association between PASK and WDR5 is strengthened at the onset of the myogenesis program, perhaps due to increased PASK nuclear translocation, as WDR5 has been reported to be localized primarily in the nucleus (Guarnaccia et al., 2021; Kikani *et al*., 2016). To expand our understanding of the biochemical interaction between WDR5 and PASK, we searched for a canonical WDR5 interaction motif (WIN) sequence within PASK exhibiting a high degree of sequence similarity with other WIN motif-containing proteins (Figure 7A).

**Figure 7.**
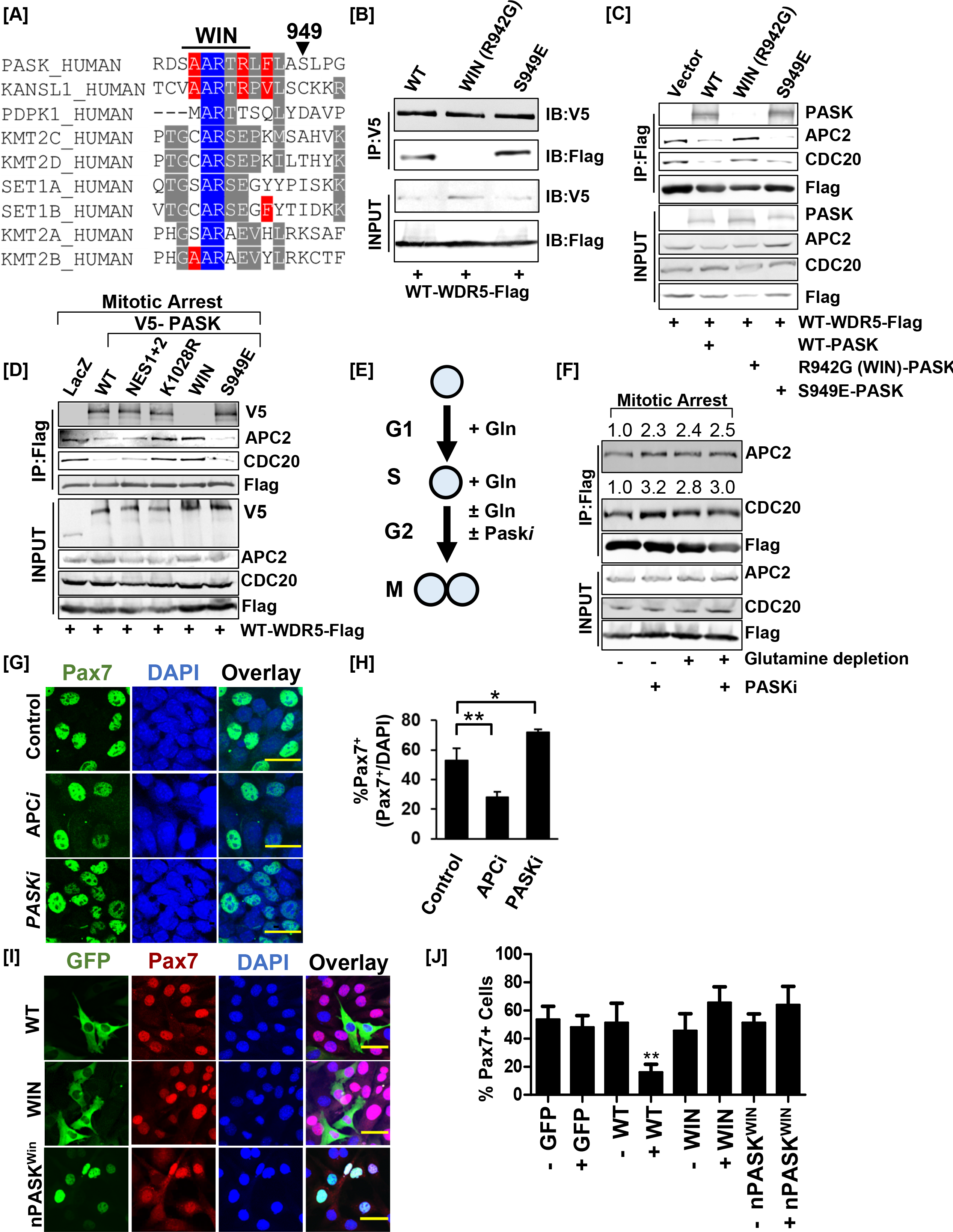
– Nuclear PASK disrupts the Wdr5-APC/C interaction causing exit from self-renewal. (A) Alignment of the hPASK WIN motif with previously established WIN motifs. S949 represents a previously reported mTOR phosphorylation site on Pask. (B) Biochemical characterization of Pask-Wdr5 interaction via R942 (WIN motif). HEK-293T cells were co-transfected with Flag-WDR5 and full-length V5-tagged WT-PASK (WT), R942G (WIN), or a phosphomimetic mutant of S949, S949E (S949E). Cells were lysed in native lysis buffer and Flag-Wdr5 was immunoprecipitated from cell extract. Co-precipitated PASK versions were detected by probing with anti-V5 antibodies. (C) PASK disrupts the WDR5-APC/C interaction. HEK-293T cells were transfected with full-length V5-tagged WT (WT), R942G (WIN), or S949E PASK, along with FLAG-WDR5. Asynchronous cells were lysed in native lysis buffer and WDR5 complexes were purified using anti-Flag beads. The immunoprecipitants were probed with PASK, APC2, CDC20, and FLAG antibodies. (D) PASK catalytic activity, phosphorylation of PASK by mTORC1, and WIN motif interaction converge to disrupt the WDR5-APC/C interaction during mitosis. HEK-293T cells expressing indicated plasmids were synchronized to the G2/M boundary by nocodazole treatment as described in methods. G2/M arrested cells were collected via centrifugation and cell pellets were lysed in native lysis buffer. Flag-WDR5 containing protein complexes were purified using anti-Flag beads. Relative enrichment of APC2, CDC20, and PASK was determined by western blotting. (E) Graphical representation of experimental scheme testing effect of glutamine withdrawal on WDR5-APC/C interaction. Cells were arrested at the G1/S boundary in the presence of glutamine. G1/S arrested cells were released into media containing 2 µM nocodazole in the presence or absence of glutamine with or without 50 µM Paski. (F) Western blot analysis of WDR5-APC/C interaction in G2/M arrested cells in (E). The numbers indicate the normalized intensities of western blot signal relative to Flag signal from IP. (G) Primary myoblasts were treated with DMSO (control), proTAME (APC/C inhibitor, APCi), or 50 µM Paski for 30hrs. Pax7+ myoblasts numbers were quantified by immunofluorescence microscopy using anti-Pax7 antibody. Scale Bar = 40 µm. (H) Quantification of % Pax7+ cells from the experiment in (G). (I) PASK-WDR5 interaction regulates myoblast identity. GFP-hPASK, GFP-hPASK^R942G^(WIN) or NLS-hPASK^L403L405S+L666S-L671A^-WIN ^R942G^(nPASK-WIN) were expressed in Pask silenced C2C12 cells. 48 hrs after retroviral infection, cells were fixed and the proportion of Pax7^hi^ cells was quantified by immunofluorescence microscopy using anti-Pax7 antibody. Scale Bar = 40 µm. (J) Quantification of % Pax7^hi^ cells from experiment in (I). \*\**P<0.005* (WT+ vs. WIN+).

The multiple alignments identified R942 as a conserved residue within the WIN motif that interacts directly with WDR5 (Figure 7A). Mutation of Arg^942^Gly (R942G) within the WIN motif ablated the interaction between PASK and WDR5 (Figure 7B). We have previously reported that mTOR phosphorylates PASK at multiple sites to promote the PASK-WDR5 association (Kikani *et al*., 2019). Three of these sites (Ser949, Ser953, Ser956) juxtapose with the WIN motif. Phosphomimetic mutation of a single mTOR phosphorylation site, Ser949 to glutamic acid (E), resulted in a stronger association between PASK and WDR5 (Figure 7B), further confirming that nutrient signaling modulates the PASK- WDR5 interaction (Kikani *et al*., 2019).

WDR5-WIN motif interaction has emerged as a key regulator of many aspects of cellular response and behavior (Guarnaccia *et al*., 2021). In stem cells, WDR5 interacts with the cell cycle linked E3 ubiquitin ligase, anaphase-promoting complex/cyclosome (APC/C) via a WIN motif. This interaction is strengthened during mitosis and is required for the rapid transcriptional reactivation of self-renewal and lineage commitment genes following mitosis in embryonic stem cells (Oh *et al*., 2020). Disruption of the WDR5-APC/C interaction reduces stem cell pluripotency. Yet, the upstream signal that regulates the WDR5-APC/C interaction remains unknown.

We have noticed rapid nuclear entry of human PASK during mitosis, prior to nuclear membrane breakdown (Figure S7A). Thus, as an upstream regulator of WDR5, we asked if PASK disrupts the WDR5-APC/C interaction during mitosis to suppress Pax7 expression. Since APC2 was demonstrated to be the only APC/C complex member that binds directly with WDR5 when the proteins are purified *in vitro* (Oh *et al*., 2020), we focused our co-immunoprecipitation experiments on APC2 and CDC20, two members of the endogenous APC/C complex that interact with WDR5 during mitosis. The presence of WT-PASK significantly disrupted WDR5-APC/C, as indicated by reduced co-precipitation of Wdr5 with APC2 and CDC20, compared with control cells (Figure 7C). PASK^WIN^ mutant, on the other hand, significantly strengthened the WDR5-APC/C interaction, suggesting that interaction of PASK with WDR5 disrupts the WDR5-APC/C interaction (Figure 7C). Since the WDR5-APC/C interaction is specifically strengthened during mitosis, we studied the role of various PASK mutants, including a catalytically dead version (KD., K1028R), in regulating the WDR5-APC/C interaction during mitosis. As shown in Figure 7D, co-expression of WT-PASK with WDR5 significantly reduced the WDR5-APC/C interaction. However, co-expression of either kinase-dead (K1028R) or WIN mutant (R942G) PASK resulted in significantly strengthened WDR5-APC/C interaction when compared to expression of WT PASK (Figure 7D). Expression of PASK^S949E^, on the other hand, further reduced the WDR5-APC/C interaction compared with WT-PASK, consistent with the stronger PASK-WDR5 association and increased PASK activity resulting from phosphorylation at the Ser949 site (Figure 7D). We previously showed that PASK phosphorylated WDR5 at S49 and that the phosphomimetic mutant of WDR5, S49E, dramatically reduced Pax7 expression and induced precocious myogenesis under proliferating conditions (Kikani *et al*., 2016). Interestingly, we find that WDR5^S49E^ completely disrupted the WDR5-APC/C interaction during mitosis, suggesting that the catalytic function of PASK regulates the WDR5-APC/C interaction (Figure S7B).

Glutamine depletion suppressed proliferation and differentiation while stimulating Pax7 expression and the preservation of stemness. Therefore, our data raise the possibility that glutamine metabolism stimulates stem cell proliferation and simultaneously drives PASK nuclear translocation to provide a window of opportunity for stem cells to disrupt the WDR5-APC/C interaction. To test this, we allowed cells to be synchronized to the G1-S boundary in media containing glutamine since the WDR5- APC/C interaction is specifically strengthened during mitosis. We then released and arrested cells at the G2/M boundary in media containing or lacking glutamine supplementation in the presence or absence of PASKi (Figure 7E). This allowed for G2/M phase-specific analysis of the role of glutamine metabolism in modulating the WDR5-APC/C interaction in stem cells. As shown in Figure 7F, PASKi treatment strengthened the WDR5-APC/C interaction in glutamine-containing media during mitosis. Interestingly, glutamine withdrawal during mitosis was sufficient to strengthen the WDR5-APC/C interaction, and the addition of PASKi in the glutamine depleted condition did not have any further additive effects (Figure 7F). This result suggests that the WDR5-APC/C interaction is a target of glutamine metabolism during mitosis in stem cells and that PASK is a mediator of glutamine signaling to WDR5-APC/C.

Cells that asymmetrically retain PASK in the nucleus after exiting cytokinesis exhibit loss of Pax7 expression in proliferating myoblasts (Figure 2G). Heterogeneity in the nuclear expression of PASK underlies heterogeneity in Pax7 expression (Figure 2E-F), which can be reversed by catalytic inhibition of PASK (Figure 6A-B). Furthermore, inhibition of glutamine metabolism reduces proliferation, inhibits PASK nuclear translocation, strengthens WDR5-APC/C interaction, and reduces heterogeneity in Pax7 expression. Therefore, we asked if APC/C activity underlies the reestablishment of Pax7 transcription after mitosis in isolated primary myoblasts. Strikingly, APC/C inhibition using the specific inhibitor proTAME resulted in a significant reduction in the number of Pax7+ cells after at least one round of the cell cycle (30 hours of treatment) compared with control cells (Figure 7G-H). Within this timeframe, PASKi treatment resulted in a significant increase in Pax7+ cell number (Figure 7G-H). Furthermore, we noticed that WT-PASK effectively downregulated Pax7 expression (Figure 7I-J) while mutation of WIN (R942G), which abolishes the PASK-WDR5 interaction, resulted in a near-complete failure in suppressing Pax7 expression with both WT and nuclear versions of PASK (Figure 7I). Finally, inhibition of APC/C resulted in the transcriptional induction of the committed progenitor and differentiation markers, *Myog* and *Mylpf*, respectively (Figure S7C). In contrast, inhibition of WDR5 or PASK, alone or in combination with APC/C inhibitor, prevented the establishment of the committed progenitor population and blocked differentiation (Figure 7G-H, S7C). These results suggest multifaceted functions of PASK and WDR5 that act in conjunction with APC/C to regulate the generation of committed progenitors and indicate that proper temporal transition of cell identity is required for successful terminal differentiation. Therefore, we discover a novel signaling role of PASK in connecting glutamine metabolism with WDR5-APC/C interaction to control stem cell fate. Our results suggest that the proliferative functions of glutamine metabolism are co-opted by stem cells to establish stem cell fate in response to the changing metabolic landscape.

## Discussion

Comprehensive studies represented in this manuscript describe a novel cell cycle-linked mechanism for establishing cell identity. Our data support a model where glutamine stimulates the proliferation of activated myoblasts to generate the committed progenitor pool during early regeneration. For these functions, glutamine metabolism drives p300/CBP-dependent acetylation of PASK. Nuclear PASK associates with WDR5 to disrupt the WDR5-APC/C interaction in self-renewing stem cells during mitotic exit. This drives exit from the self-renewal program to generate committed progenitors primed for differentiation. The *in vivo* relevance of our data is evident from our observation that either genetic loss of PASK (Figure 6C-D), or pharmacological inhibition of glutamine metabolism during muscle regeneration resulted in increased Pax7+ MuSC numbers (Figure 5H-I). Curiously, PASK expression declines by 7 days post muscle injury, which is also the timepoint when the Pax7+ self-renewing population is reestablished at the end of early myogenesis (Cutler *et al*., 2021). Thus, as a metabolic sensory kinase enriched in stem cells, PASK functions to integrate metabolic (via glutamine) and nutrient signaling (via mTOR) inputs with cell cycle control to orchestrate exit from self-renewal and promote the onset of the terminal differentiation program.

PASK is critically required for *in vitro* differentiation and *in vivo* muscle regeneration (Kikani *et al*., 2016). Biochemically, PASK is a constitutively active kinase in uninduced conditions, unlike typical activation-loop activated protein kinases. PASK is highly expressed in self-renewing stem cells, yet, inhibiting PASK preserves stemness (Figure 1). Our data presented here resolve this incongruency by proposing that rapid nucleocytoplasmic shuttling of PASK modulates the expression of stemness-linked genes. Differentiation signals, including glutamine and mTOR, stimulate the activation and persistent nuclear localization of PASK, where it interacts with WDR5 to drive exit from self-renewal and differentiation. Therefore, nucleo-cytoplasmic shuttling enables PASK to precisely compartmentalize its functions between permitting self-renewal and driving differentiation in response to the appropriate nutrient signals. In agreement, our results suggest that nuclear accumulation of PASK is signal-regulated via p300/CBP-induced acetylation and that mitochondrial glutamine metabolism is a primary source of acetyl-CoA for driving both PASK acetylation and nuclear translocation.

Inhibition of PASK with PASKi also increased Pax7^hi^ cell numbers. Furthermore, *PASK*^KO^ animals exhibit increased Pax7+ cell numbers during regeneration at day 5 and day 14. In the context of muscle regeneration, these results are significant. In WT mice, PASK expression begins to decline by Day 7, and by Day 15 post-injury, its expression is nearly completely lost. This timepoint of PASK expression decline coincides with the emergence of the self-renewing population during regeneration, according to a recent study (Cutler *et al*., 2021). Thus, the downregulation of PASK expression could initiate the reactivation of Pax7 expression needed for the self-renewal program following the initial burst of regenerative myogenesis.

Taken together, our studies provide a novel model for connecting the metabolic and nutrient signaling environment with a mechanistic cell cycle-associated control of stem cell fate. Our parallel results in mouse ES cells further suggest the generalizability of PASK function to stem cells of varied tissue origins. The downstream target of PASK function, the WDR5-APC/C interaction, is a conserved feature among both mESCs and muscle stem cells. Transient inhibition of PASK in both mESCs and MuSCs preserves cell-identity, self-renewal and stemness. Thus, as a druggable, stem cell specific kinase, PASK represents an excellent candidate for future therapeutic applications where the maintenance of stem cell function is desired.

## Methods

### Key Resources Table

| Antibodies |  |  |
| --- | --- | --- |
| REAGENT | SOURCE | IDENTIFIER |
| Anti-Pax7 | DSHB, University of Iowa | PAX7 |
| Anti-MyoG | DSHB, University of Iowa | F5D |
| Anti-MHC | DSHB, University of Iowa | MF 20 |
| Anti-Laminin | Sigma Aldrich | L9393 |
| Anti-FLAG | Sigma Aldrich | F3165 |
| Anti-V5 | ThermoFisher | R960-25 |
| Anti-PASK | Cell Signaling Technology | C70B2 |
| Anti-Acetylated lysine | Cell Signaling Technology | 9814 |
| Anti-APC2 | Cell Signaling Technology | 12301 |
| Anti-CDC20 | Cell Signaling Technology | 14866 |
| Anti-$\alpha/\beta$-Tubulin | Cell Signaling Technology | 2148 |
| Anti-KAT2A (GCN5L2) | Cell Signaling Technology | 3305 |
| Anti-BrdU | ThermoFisher Scientific | PA5-32256 |
| TruSeq Stranded Total RNA<br>Library Prep Kit with Ribo-Zero<br>Gold | Illumina | RS-122-2302 |
| Cell Lines |  |  |
| HEK 293T | ATCC |  |
| C2C12 | ATCC |  |
| Mouse ES Cells | ATCC |  |
| Primary myoblasts | Isolated fresh from mice |  |
| Chemicals |  |  |
| **REAGENT** | **SOURCE** | **IDENTIFIER** |
| BPTES | Selleck Chem | S7753 |
| A-485 | Selleck Chem | S8740 |
| UK5099 | Selleck Chem | S5317 |
| BioE-1197 | Jared Rutter, University of Utah | Not applicable |
| Rapamycin | LC Labs | R-5000 |
| Akt inhibitor IV | Sigma Aldrich | 124011 |
| LY294002 | Cell Signaling Technology | 9901 |

### Resource Availability

Further information and requests for reagents and resources should be directed to the lead contact, Chintan Kikani.

### Materials Availability

Plasmids generated in this work are available upon request. Please direct inquiries to the lead contact. Other reagents used are commercially available.

### Data and Code Availability

RNA sequencing gene expression data from C2C12 myoblasts treated with Paski will be uploaded upon successful review.

### Experimental Model and Subject Details

#### Mouse models

WT and *Pask^-/-^* animals were described previously (Kikani *et al*., 2016). Animal experiments were performed in accordance with protocols approved by the Institutional Animal Care and Use Committee at the University of Kentucky to CK (2019-3317). Animals were housed in the environment in compliance with the Guide for the Care and Use of Laboratory Animals in an AAALAC-accredited facility at the University of Kentucky on a 12:12-hour light: dark cycle.

#### Cell Culture

HEK 293T and C2C12 myoblasts were acquired from the American Type Culture Collection (ATCC). Cells were cultured in DMEM (Gibco) with 10 % fetal bovine serum (FBS) and 1 % penicillin/streptomycin (pen-strep) at 37 °C with 5 % CO2. Cells were passaged every two days. For glutamine and serum withdrawal experiments, cells were washed once with 1 X PBS and cultured for 12 hours in DMEM lacking glutamine or serum (Gibco), before re-stimulation with glutamine or serum for the time period indicated. For differentiation experiments, DMEM containing 2 % horse serum and 1 % pen-strep was used to induce differentiation of C2C12 mouse myoblasts. Differentiation medium (DMEM with 2 % horse serum) was refreshed every 24 hours. For stable expression in 293T and C2C12 cells, puromycin selection proceeded for two weeks at a concentration of 2 µg/mL. Cells were counted using the Countess 3 Automated Cell Counter (ThermoFisher).

#### mESC culture

mESCs were previously generated from C57BL/6 x 129 F1 male embryos (Banaszynski et al., 2013). mESCs were maintained on gelatin-coated plates in 2i/LIF medium containing a 1:1 mix of DMEM/F-12 (Gibco) and Neurobasal medium (Gibco) including N-2 supplement (Gibco), B-27 supplement (Gibco), 2-mercaptoethanol, 2 mM Glutamax (Gibco), LIF (produced in house) supplemented with 3 µM CHIR99021 (Stemgent) and 1 µM PD0325901 (Stemgent) (2i). When assessing the effect of PASKi on mESCs, the 2i supplement was replaced with 75 µM of BioE-1197 + LIF (PASKi).

#### EB formation assay

For EB formation, mESCs were diluted to 10^4^ cells/ml in EB differentiation media (DMEM, 15 % FBS, 1x MEM-NEAA, 1x Pen/Strep, 50 μM β-mercaptoethanol) and 30 μl drops were placed on the lid of a 150 mm dish. The lid was inverted and placed over a dish containing 10–15 ml of PBS. The hanging drops were cultured for 3 days at 37 ℃ and 5 % CO2. The hanging drops were then washed from the lids with EB differentiation media and cultured in 100 mm dishes on an orbital shaker at 50 rpm for an additional day.

### Methods

#### Preparation of Protein Lysates

293T cells or C2C12 cells were seeded in 100 mm dishes overnight prior to transfection using 1:3 ratio of DNA to JetPrime reagent (Polyplus) in JetPrime Buffer (Polyplus) at approximately 80 % confluency, according to manufacturer protocol. Cells were scraped and lysed 24 hours following transfection in native cell lysis buffer (20 mM Tris-HCl pH 7.5, 150 mM NaCl, 1 mM EDTA, 1 mM NaF, 1 mM beta-glycerophosphate, 1% Triton X-100) containing freshly added protease and phosphatase inhibitors (1 mM PMSF, 1X protease inhibitor cocktail, 1 X phosphatase inhibitor cocktail; Sigma) for 20 minutes on ice. For denaturing lysis to prepare samples for acetyl-lysine pulldown, homogenized tissue samples or cells were lysed in RIPA lysis buffer (25 mM Tris pH 7.6, 150 mM NaCl, 1 % NP-40, 1 % sodium deoxycholate, 0.1 % SDS) containing freshly added protease and phosphatase inhibitors (1 mM PMSF, 1X protease inhibitor cocktail, 1 X phosphatase inhibitor cocktail; Sigma). Whole-cell lysates were cleared by centrifugation at 15,000 rpm for 20 minutes. Protein quantification was performed using BCA kit (Pierce), following manufacturer instructions.

#### Co-Immunoprecipitation

Cell lysates were prepared as described above. 10 % of each lysate was collected (input) and the remaining lysate was incubated with Anti-V5 Agarose Affinity Gel beads (Sigma) or Anti-FLAG M2 magnetic beads (Sigma) overnight at 4 °C. Immunoprecipitation (IP) samples were washed the following day with wash buffer (20 mM Tris-HCl pH 7.5, 150 mM NaCl, 1% Triton X-100) on ice for five times each. For acetyl-lysine pulldowns, IP samples were washed with RIPA lysis buffer (25 mM Tris pH 7.6, 150 mM NaCl, 1 % NP-40, 1 % sodium deoxycholate, 0.1 % SDS). Samples were resolved via SDS-PAGE and immunoblot analysis was conducted with the antibodies listed.

#### Subcellular Fractionation

For subcellular fractionation, C2C12 cells were seeded in 100 mm dishes and scraped directly into phosphate buffer saline (PBS). Cells were subsequently centrifuged at 1000 rpm for 2 minutes at 4 °C. The pellet was washed with additional PBS twice (1000 rpm for 2 minutes for each wash), before the pellet was resuspended in 200 uL of ice-cold fractionation buffer (10 mM HEPES, 10 mM KCl, 1.5 mM MgCl2, 0.34 mM sucrose, 10 % glycerol, 1 mM DTT, 1 mM PMSF, protease inhibitor cocktail). Triton X-100 was subsequently added to 0.1 % final concentration. The lysate was incubated on ice for 8 minutes, before centrifugation at 1,300 x g for 5 minutes at 4 °C. The supernatant fraction was collected, while the pellet washed twice in additional fractionation buffer containing 0.1 % triton. The collected supernatant fraction was centrifuged at 20,000 x g for 5 minutes at 4 °C (collecting the supernatant as the cytosolic fraction). The pellet (nuclear fraction) was resuspended in lysis buffer containing 20 mM HEPES pH 7.9, 1.5 mM MgCl2, 0.5 M NaCl, 0.2 mM EDTA, 20 % glycerol, 1 % Triton X-100, 1 mM PMSF, protease inhibitor cocktail for 10 minutes at 4 °C. The nuclear fraction was clarified via centrifugation at 20,000 x g for 5 minutes at 4 °C.

#### Isolation of primary myoblasts

MuSCs were isolated from 10–12-week old C57BL6 mice according to published protocols (Danoviz and Yablonka-Reuveni, 2012). Briefly, TA muscles from hind-limbs of mice were isolated, minced in DMEM and enzymatically digested with 0.1% Pronase for 1 hr. After repeated trituration, the cell suspension was filtered through a 100 μM filter. Cells were plated on Matrigel-precoated plates in myoblast growth media (DMEM + 20% FBS + 1% Chicken Embryo Extract (CEE) + 1ng/ml b-FGF) in the presence or absence of 25 µM BioE-1197 or 2 µM BPTES. For the glutamine depletion experiment, growth media replaced with DMEM lacking glutamine (ThermoFisher Inc, Cat#11960044) containing 20% FBS + 1%CEE + 1ng/ml b- FGF.

#### Cell synchronization

Cell synchronization was performed as described previously with minor modification (Kikani et al., 2012). For cell synchronization, cells were plated at 5 × 10^5^ cells per 100 mm dish containing DMEM supplemented with serum and antibiotics. After 24 hours, growth media was replaced with media containing 5% serum with aphidicolin (2 μg/ml) for 12 hours, and then with media containing 1% serum with aphidicolin (2 μg/ml) for 8 hours to synchronize the cells in the G1 phase. To arrest cells in G2/M phase, cells were released in media containing DMEM + 10% FBS + 1% PS + 2 µM nocodazole for 24 hours. M-Phase arrested cells were collected by trituration followed by centrifugation at 1500 RPM X 5 mins at 4C. Cell pellets were prepared as described above to perform immunoprecipitation experiments. For testing the effect of glutamine in mitotic cells, aphidicolin arrested cells were washed twice with media lacking glutamine prior to releasing cells in DMEM without glutamine + 10% FBS + 1% PS + 2 µM nocodazole.

#### SDS-PAGE and Immunoblotting

Whole cell lysates (15 µg) or IP samples were denatured at 95 °C for five minutes before they were loaded into 10 % SDS-PAGE gels submerged in running buffer (25 mM Tris, 192 mM glycine, 0.1 % SDS). A constant voltage of 130 V was applied for 1.5 hours. Protein samples were transferred to nitrocellulose membranes (0.45 um) for 1.5 hours at 100 V (constant voltage) in 1X Towbin Buffer (25 mM Tris, 192 mM glycine, 20 % (v/v) methanol, 0.05 % sodium dodecyl sulfate pH 8.3). Membranes were subsequently blocked in 3 % fish gelatin (Sigma) dissolved in TBS-T (20 mM Tris pH 7.5, 150 mM NaCl, 0.1 % Tween-20) for one hour prior to incubation with the primary antibodies specified overnight at 4 °C. Following incubation, membranes were washed three times with TBS-T prior to immunoblotting with the HRP-linked or fluorescent secondary antibodies for one hour at room temperature. Blots were then washed three times with TBS-T and incubated with enhanced chemiluminescent (ECL) substrate (Clarity, Biorad) for one minute to detect HRP-linked antibodies or washed in PBS for five minutes (for fluorescent antibody detection). ChemiDoc MP (Biorad) was used for ECL detection and Odyssey CLX (LI-COR) was used for fluorescent detection at near-IR wavelengths of 680 nm or 800 nm.

#### Immunofluorescence and Immunohistochemistry

For all immunofluorescence microscopy, cells were seeded in 24 well plates on glass coverslips pre-coated with 0.1% gelatin (HEK293T or C2C12 cells) or 1 mg/ml Matrigel. At the end of all experimental timepoints, cells were fixed with 4% paraformaldehyde (EM grade, Electron Microscopy Services) in 1 X PBS for 15 minutes, before wells were washed three times with 1 X PBS. Samples were subsequently permeabilized with 0.2 % Triton X-100 in 1 X PBS for 10 minutes at room temperature, before blocking in 10 % normal goat serum in 1 X PBS for one hour. Primary antibodies indicated were diluted in 1 X PBS and subsequently added to all wells. Plates were incubated overnight in a humidified chamber at 4 °C, before wells were washed three times with 1 X PBS. Secondary antibodies (Alexa Fluor, Thermo Fisher) were then added to the wells and incubation proceeded for one hour at room temperature. The wells were subsequently washed three times with 1 X PBS, before coverslips were mounted with ProLong Diamond Antifade mounting media with DAPI (Thermo Fisher) onto glass slides. After coverslips were allowed to cure overnight at room temperature, the slides were imaged with confocal microscopy (Nikon A1R). Images were analyzed and quantified using Fiji software. For quantification of microscopic images of cells, at least 5 separate fields were counted for each experiment with at least 25 cells counted per field.

#### Immunohistochemistry for Pax7 staining

For detection of Pax7 from the tissue sections, freshly isolated muscles at indicated timepoints after muscle injury were fresh-frozen in OCT in 2-methylbutane. Muscles were cross-sectioned at 10 μm, air-dried for 20 mins and fixed for 20 minutes in 4% paraformaldehyde (PFA) in PBS. Sections were subjected to antigen retrieval using a 2100 PickCell Retriever followed by quenching for 5 minutes in 3% H2O2. Tissue sections 60 minutes in 5% normal goat seum, followed by mouse on mouse block for 30 mins. Sections were incubated overnight at 4°C in appropriate primary antibody, washed 3 times in PBS and incubated for 2 hours at room temperature in secondary antibody. Sections were washed in PBS and Tyramide signal amplification (Biotium) was performed according to manufacturer’s instructions. A glass coverslip was mounted using Prolong Anti-Fade reagent with DAPI.

#### RNASeq Methods and Analysis

Total RNA was isolated from cultured C2C12 myoblasts using an RNeasy Mini Kit (QIAGEN). Agilent 2100 TapeStation was used to determine RNA quality and samples with RNA Integrity Numbers (RIN) 8 were chosen for RNA seq . Libraries were prepared using TruSeq Stranded Total RNA Library Prep Kit with Ribo-Zero Gold (Illumina). Sequencing was performed on the Illumina Hiseq 2500 with Paired-End reads. De-multiplexed read sequences were aligned to the *Mus musculus* mm10 reference sequence using TopHat2 splice junction mapper. Raw counts were normalized, and differentially expressed genes were called using DESeq2 analysis. Heatmaps, volcano plots, and hallmark pathway analysis were generated by R. Gene set enrichment analysis (GSEA, (Mootha et al., 2003; Subramanian et al., 2005)) was performed using the hallmark gene sets (MSigDB hallmark gene sets) and a previously identified gene signature for stemness (Wong et al., 2008).

#### Quantitative Real-time PCR

Total RNA was extracted using the RNeasy Kit (Qiagen) after all treatments are accomplished according to the protocol supplied by the manufacturer. RNA was quantified using a Implen MP80 and 1 µg of RNA was used for reverse transcription (QuantiTect Reverse Transcription Kit). cDNA samples were diluted 1:16 and used for real-time qPCR (Applied Biosystems) using Power Up SYBR Green PCR Master Mix (Applied Biosystems). Standard curve was prepared to obtain relative quantity for each experimental primer sets which were normalized with 18s rRNA levels.

#### Muscle Injury and Regeneration

Muscle Injury and Regeneration Animal experiments were performed in accordance with protocols approved by the Institutional Animal Care and Use Committee at the University of Kentucky to CK (2019- 3317). For muscle injury, 1.2% BaCl2 was freshly prepared in sterile distilled water, and 25 µl was injected intramuscularly into TA muscles of anesthetized mice. For BPTES or DMSO injection, 10µM working stock of BPTES was prepared in sterile 0.1X PBS. 10µl of 10µM BPTES or 0.001% (v/v) DMSO in 0.1X PBS was intramuscularly injected into TA muscles 5 days after the BaCl2 induced injury every 24 hours for 3 days. Animals were monitored to ensure full recovery from anesthesia and followed for a duration set forth by experimentation.

#### Metabolomics Sample Preparation

3 x 10^5^ cells/well with 5 replicates per condition were seeded in 6-well plates one day before metabolite extraction. An identical cell counting plate was seeded for each condition. Cell culture media was aspirated thoroughly from cell culture wells and rapidly washed twice with 10 mL and 5 mL of 0.1 X PBS and subsequently placed on ice after PBS washes. Metabolite extraction buffer was added to each well (1000 µL of 50% Methanol with 20 μM L-norvaline (as an internal control) on ice and plates were then transferred to -20 °C for 10 minutes. Cells were then scraped with a cell scraper (Sarstedt) and the entire volume of each well was transferred to 1.5 mL Eppendorf tubes on ice. Samples were then thoroughly homogenized on a disruptor genie (Scientific Industries) at 3000 rpm for 3 minutes at room temperature. After homogenization, samples were then centrifuged at 15,000 rpm for 10 minutes at 4 °C. The top 90 % of sample supernatant was transferred to 1.5 ml tubes and samples were stored at -80 °C until GC-MS metabolite quantification. Once supernatant was removed, the insoluble pellet was washed four times with 50% methanol before a final wash of 100 % methanol. Between each wash, pellets were homogenized on a disruptor genie at 3000 rpm for 1minute and then spun down at 15,000rpm at 4°C. Following washes, the pellet was hydrolyzed in 3N HCl for 2 hours at 95 °C on a shaking Thermomixer (Eppendorf). The reaction was quenched with 100 % methanol containing 40 μM L-norvaline (as an internal control). The sample was then incubated on ice for at least 30 min. The supernatant was collected by centrifugation at 15,000 rpm at 4 °C for 10 min and both polar and hydrolyzed samples subsequently dried by vacuum centrifuge at 10^-3^ mBar (Andres et al., 2020; Sun et al., 2019).

#### Metabolomics Analysis

Dried polar and hydrolyzed samples were derivatized by the addition of 20 mg/mL methoxyamine hydrochloride in pyridine and incubation for 1.5 hr at 30 °C. Sequential addition of N-methyl-trimethylsilyl-trifluoroacetamide (MSTFA) followed with an incubation time of 30 min at 37 °C with thorough mixing between addition of solvents. The mixture was then transferred to a v-shaped amber glass chromatography vial. An Agilent 7800B gas-chromatography (GC) coupled to a 7010A triple quadrupole mass spectrometry detector equipped with a high-efficiency source was used for this study.

GCMS protocols were similar to those described previously (Sun et al.; Young et al., 2020), except a modified temperature gradient was used for GC: Initial temperature was 130 °C, held for 4 min, rising at 6 °C/min to 243 °C, rising at 60 °C/min to 280 °C, held for 2 min. The electron ionization (EI) energy was set to 70 eV. Scan (*m/z*: 50 – 800) and full scan mode were used for target metabolite analysis.

Metabolite EI fragmentation pattern and retention time were determined by ultrapure standard purchased from Sigma. Ions (m/z) and retention time (min) used for glycogen quantitation was glucose (160 or 319 *m/z*; 17.4min), Relative abundance was corrected for recovery using the L-norvaline standard and adjusted to protein input represented by pooling amino acids detected by GC-MS (Andres *et al*., 2020; Sun *et al*., 2019)

### Quantification and Statistical Analysis

A two-tailed t-test was performed in GraphPad Prism 9 to determine significant differences between relevant sample and control groups (indicated in Results). *, **, *** on graphs represent p < 0.05, p < 0.01, p < 0.001, respectively. A two-tailed t-test was used to compare differences. Error bars are represented as the standard error of the mean as indicated where n ≥ 3.

## Acknowledgments

We acknowledge Jared Rutter for the critical reading of the manuscript and Jared Rutter and Wojciech Swiatek for providing a PASK inhibitor. In addition, we thank Cory Dungan (Center for Muscle Biology, University of Kentucky) and Kevin Murach (University of Arkansas Medical Center) for stimulating discussion and providing resources, reagents, and training in performing muscle histology.

## Author contributions

CKK conceived the project, CKK and MX designed the experiments. MX, CHW, GM, BK, DBC, SM, PS, and ALD performed experiments and analyses. LEAY performed metabolomics experiments. MSG, RCS, LAB, and CKK analyzed the data and interpretation. CKK, MX, and LAB wrote the manuscript with contributions and inputs from all authors.

## Supplementary Materials

### Supplementary Figure Legends

**Figure S1.**
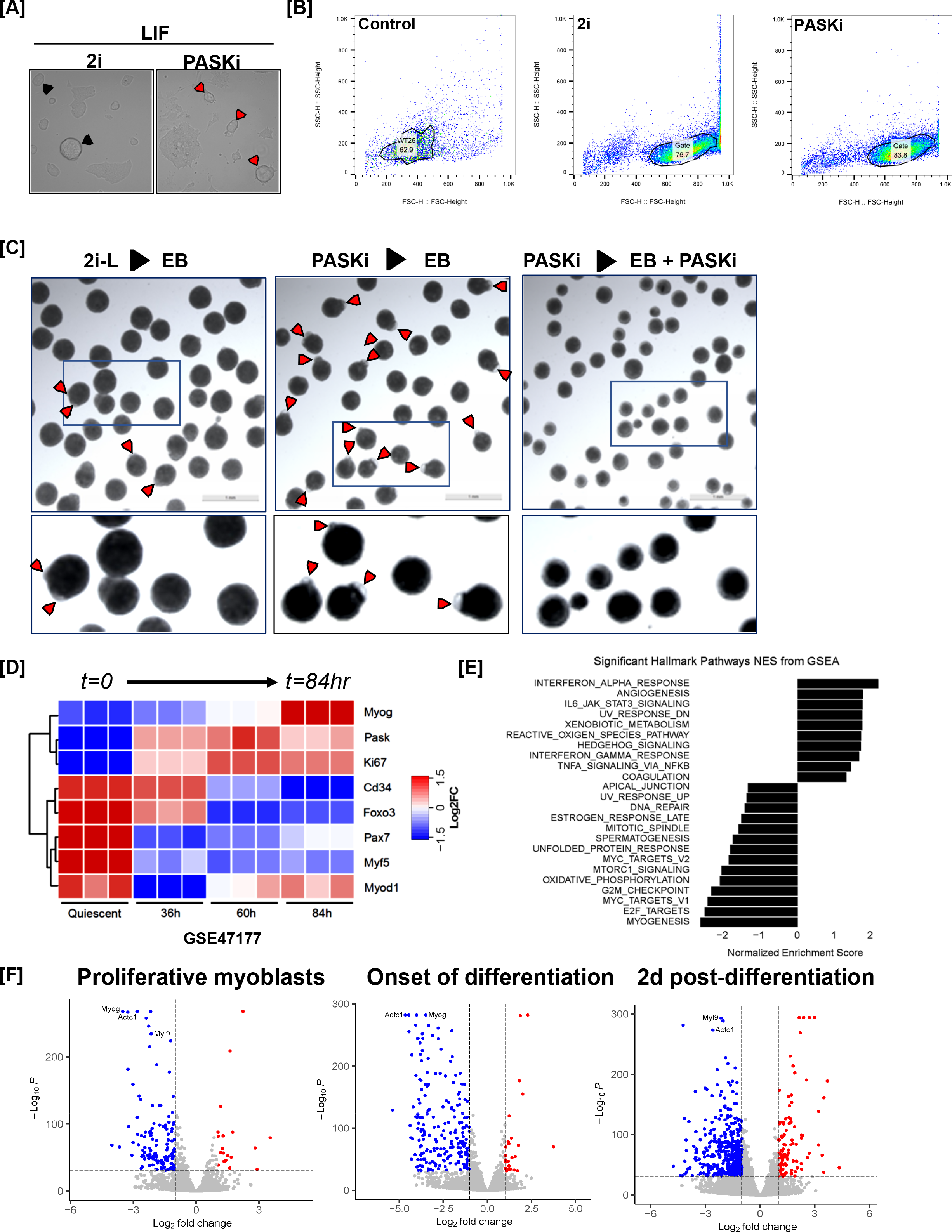
Transient PASK inhibition enhances differentiation potential while prolonged PASK inhibition prevents differentiation. (A) Morphological comparison of mESCs grown in 2i media vs 75 µM PASKi media. mESCs were first cultured in 2i media to maintain pluripotency. 2i media was subsequently changed to 2i or Paski. Cells were cultured in this media for 4 days. (B) Flow cytometry gating from an experiment in Figure 1A. (C) Embryoid body formation assay following 2i or PASKi withdrawal. Arrows indicate an embryoid body showing at least a small, translucent fluid-filled cavitation. Box indicates a magnified section of the field. Scale bar = 100 µm. (D) Heatmap of muscle stem cell associated gene expression in isolated muscle stem cells during regeneration (Gene Expression Omnibus GSE47177). (E) Significantly enriched and depleted hallmark pathways from gene set enrichment analysis (GSEA) in Paski treated versus control C2C12 myoblasts 2 days following the onset of differentiation. (F) RNA-sequencing volcano plot of differentially expressed genes in Paski treated versus control in proliferating, differentiation onset, or 2 d post-differentiation onset C2C12 myoblasts (red – significantly upregulated genes, blue – significantly downregulated genes).

**Figure S2.**
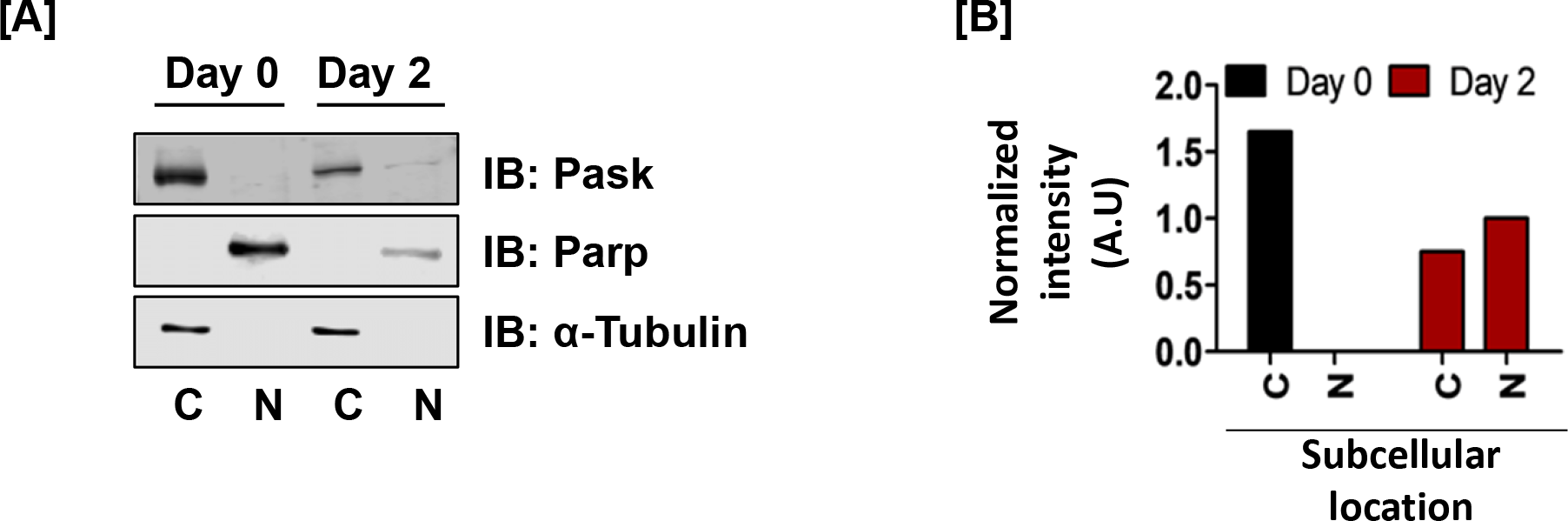
A fraction of PASK is nuclear during myoblast differentiation. (A) Cytoplasmic and nuclear fractionation of C2C12 cells at Day 0 and Day 2 showed increased Pask nuclear accumulation during differentiation. Notice reduced levels of Parp in the nuclear fraction of differentiated cells (Day 2) compared with proliferating cells (Day 1). (B) Relative abundance of Pask in cytoplasmic or nuclear fraction at Day 0 or Day 2. For cytoplasmic and nuclear fractions, tubulin and Parp, respectively, were used to quantify Pask abundance.

**Figure S3.**
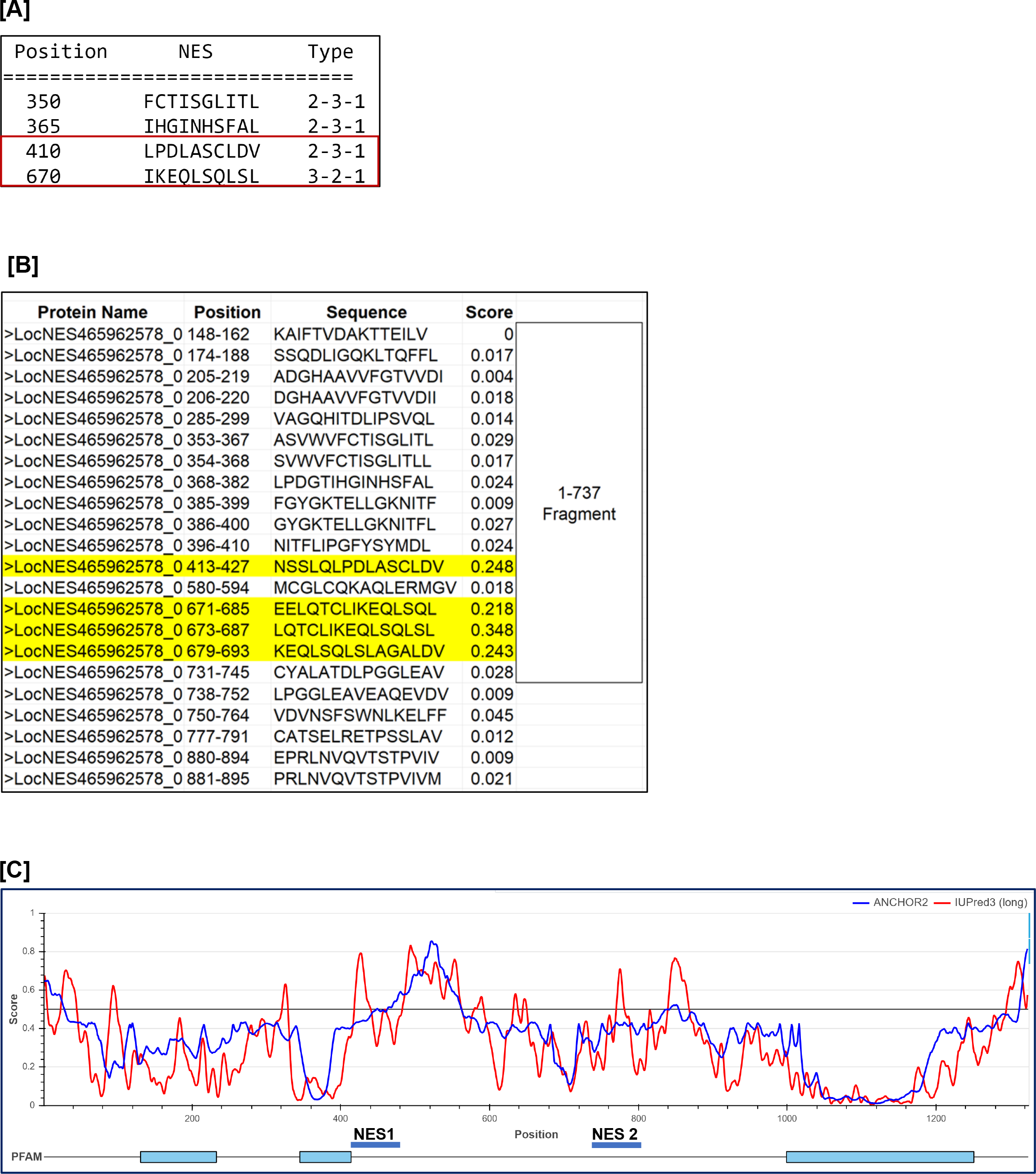
Prediction of PASK nuclear export sequences. (A) Output from NES Finder 0.2 program (Fornerod lab) indicating score for predicated nuclear export sequences in human PASK. (B) LosNES output indicating score for predicated nuclear export sequences in human PASK. Highlighted sequences shows two highest ranking NES sequences corresponding to NES1 and NES2 sequences discussed in the manuscript. (C) Organization of secondary structure of human PASK that could participate in protein-protein interaction during nuclear export. Anchor scores indicated disorder structure that is predicted to engage in protein-protein interactions.

**Figure S4.**
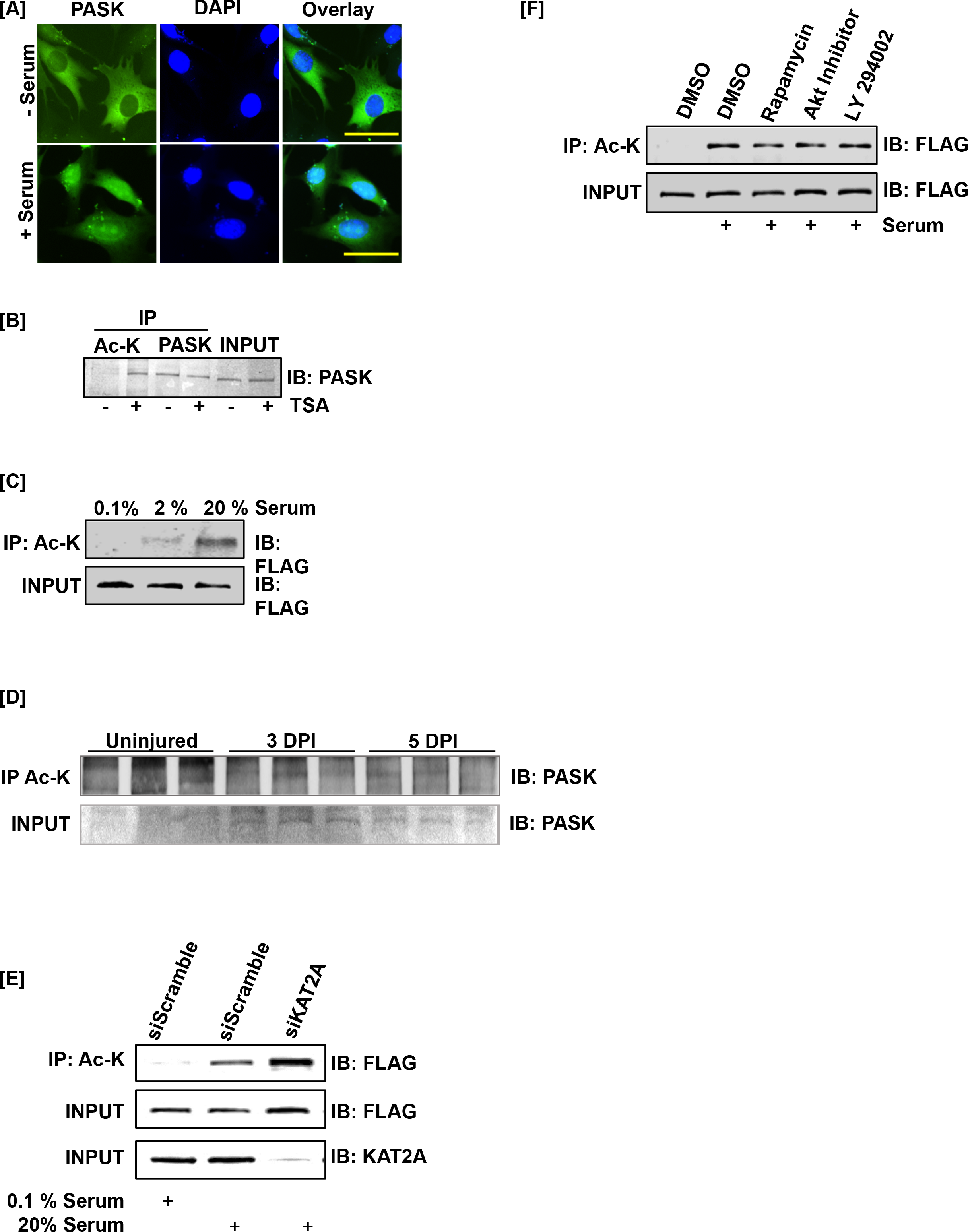
PASK is acetylated *in vitro* and *in vivo* during muscle regeneration. (A) Endogenous Pask translocated to the nucleus in response to acute serum stimulation. C2C12 cells were serum-starved overnight. Cells were stimulated with 20% serum for 2 hours where indicated, followed by immunofluorescence detection of mouse Pask. (B) Acetylation of PASK in C2C12 myoblasts. Proliferating C2C12 myoblasts were treated with trichostatin A (TSA). The lysate was immunoprecipitated with total acetyl-lysine (Ac-K) prior to immunoblotting for endogenous Pask. (C) HEK-293T cells stably expressing FLAG-tagged full-length PASK (aa 1-1323) were serum-starved (0.1 % serum) for 12 hours. Cells were subsequently stimulated with either 0.1 % serum or 20 % serum for 2 hours. The lysate was immunoprecipitated with total acetyl-lysine (Ac-K) prior to immunoblotting for FLAG. (D) HEK-293T cells stably expressing FLAG-tagged full-length PASK (aa 1-1323) were serum-starved (0.1 % serum) for 12 hours. Cells were treated with 100 nM rapamycin during the 12-hour serum withdrawal or with 1 μM Akt Inhibitor IV or 25 μM LY 294002 for one hour prior to serum stimulation. Cells were subsequently stimulated with either 0.1 % serum or 20 % serum for 2 hours. The lysate was immunoprecipitated with total acetyl-lysine (Ac-K) prior to immunoblotting for FLAG. (E) Acetylation of Pask *in vivo. Pask^+/-^* mice were injured with 1.2 % w/v BaCl2 to the tibialis anterior muscles. Tissues were harvested at day 0 (uninjured), day 3, and day 5 post-injury. Lysates were immunoprecipitated with total acetyl-lysine (Ac-K) prior to immunoblotting for endogenous Pask. (F) HEK-293T cells stably expressing FLAG-tagged full-length PASK (aa 1-1323) were transfected with scrambled or KAT2A targeting siRNAs. Cells were serum starved (0.1 % serum) for 12 hours and subsequently stimulated with either 0.1 % serum or 20 % serum for 2 hours. The lysate was immunoprecipitated with total acetyl-lysine (Ac-K) prior to immunoblotting for FLAG.

**Figure S5.**
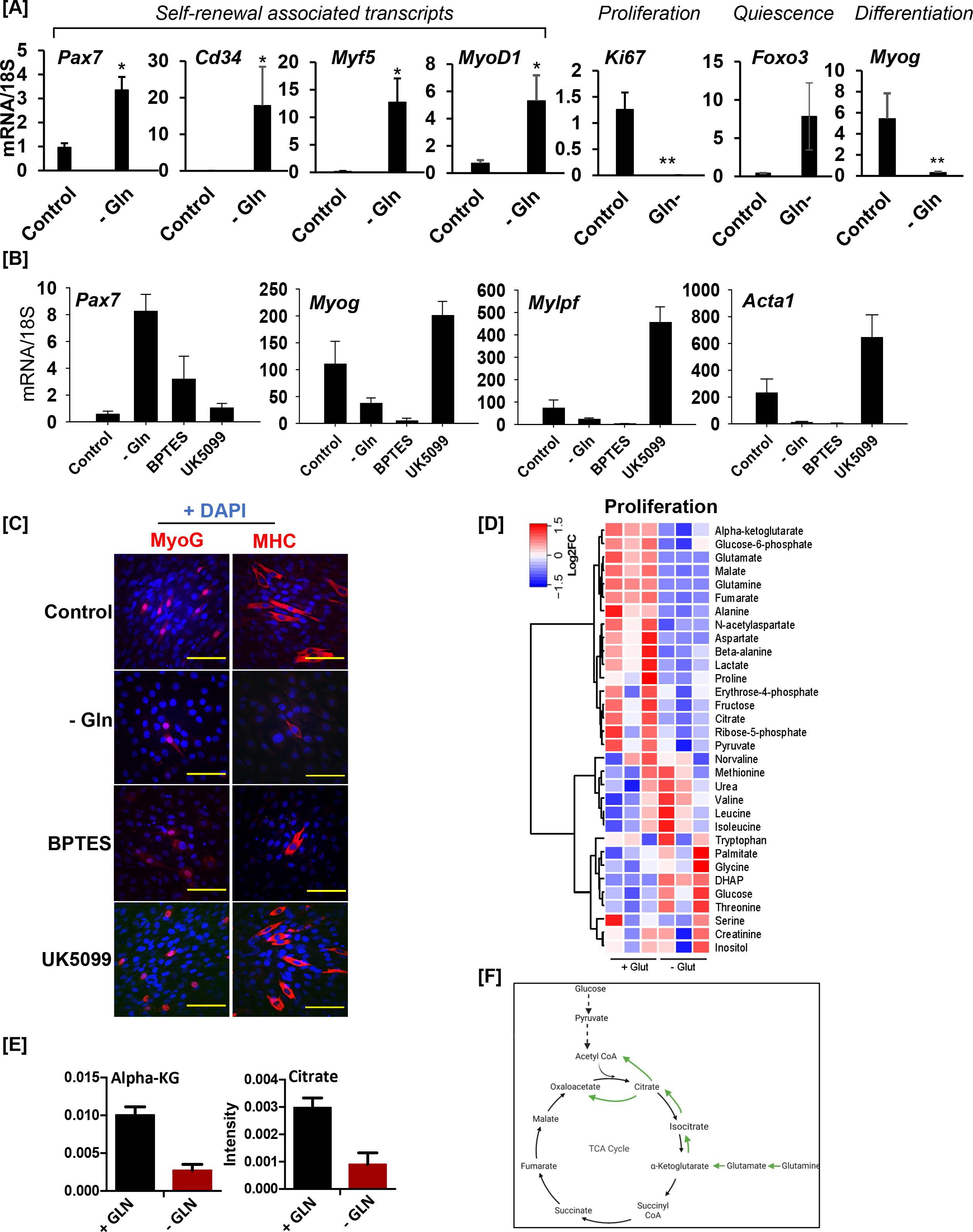
Glutamine depletion preserves stem cell identity while antagonizing differentiation. (A) Expression of genes associated with muscle stem cell self-renewal (Pax7, Cd34, Myf5, and Myod1), proliferation (*Ki67)*, quiescence (Foxo3) and differentiation (*Myog)* in primary myoblasts cultured with (Control) or without (- Gln) glutamine, as quantified by RT-qPCR. (B) Expression of indicated genes in C2C12 myoblasts cultured in the presence or absence of 2 mM glutamine or in the presence of 10 µM BPTES or 10 µM UK5099 in glutamine containing media. (C) C2C12 myoblasts were cultured in the presence (Control, BPTES, UK5099) or absence (- Gln) of glutamine and treated with BPTES (glutaminolysis inhibitor) or UK 5099 (mitochondrial pyruvate carrier inhibitor) as indicated. C2C12 myoblasts were induced to differentiate for 2 days, and cells were analyzed by immunofluorescence against Myog and MHC. Scale bar = 40µM (D) Heatmap of metabolite concentrations in differentiating (day 1) C2C12 myoblasts cultured in the presence (+ Glut) or absence (- Glut) of glutamine. (E) Levels of indicated TCA cycle metabolites (alpha-KG and Citrate) that are implicated in glutamine- derived acetyl-CoA generation for signaling and epigenetic functions. (F) Illustration of glucose and glutamine-derived metabolic pathways leading to the generation of intracellular acetyl-CoA.

**Figure S6.**
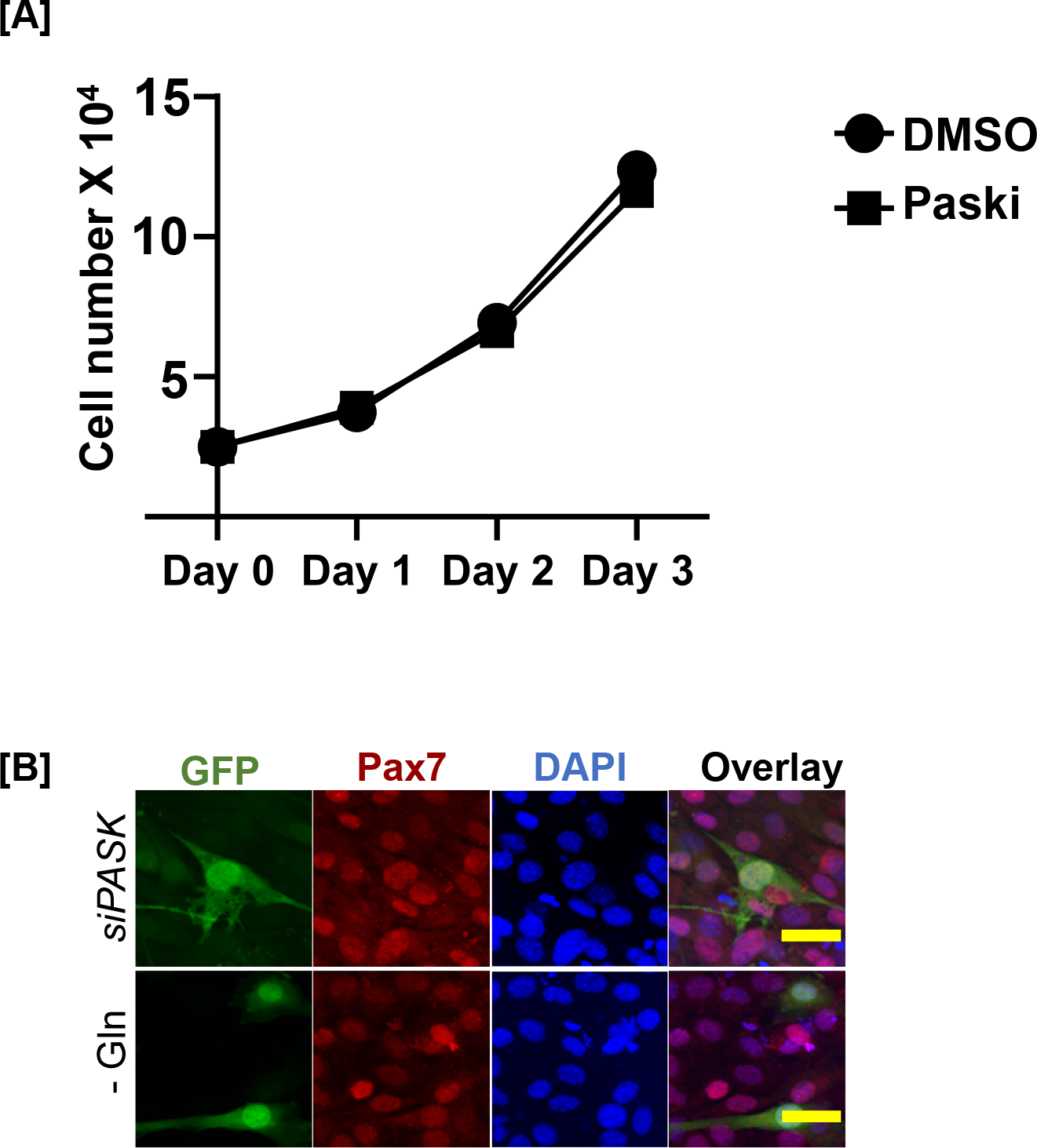
Glutamine driven generation of differentiation competent progenitors requires nuclear PASK function. (A) Growth curve of C2C12 myoblasts cultured in the presence of DMSO (vehicle control) or 50 µM Paski treatment over 3 days. (B) Expression of control vector, GFP did not reverse the observed increase in Pax7 levels in C2C12 cells due to *Pask* silencing or glutamine withdrawal. See Figure 6H for quantification.

**Figure S7.**
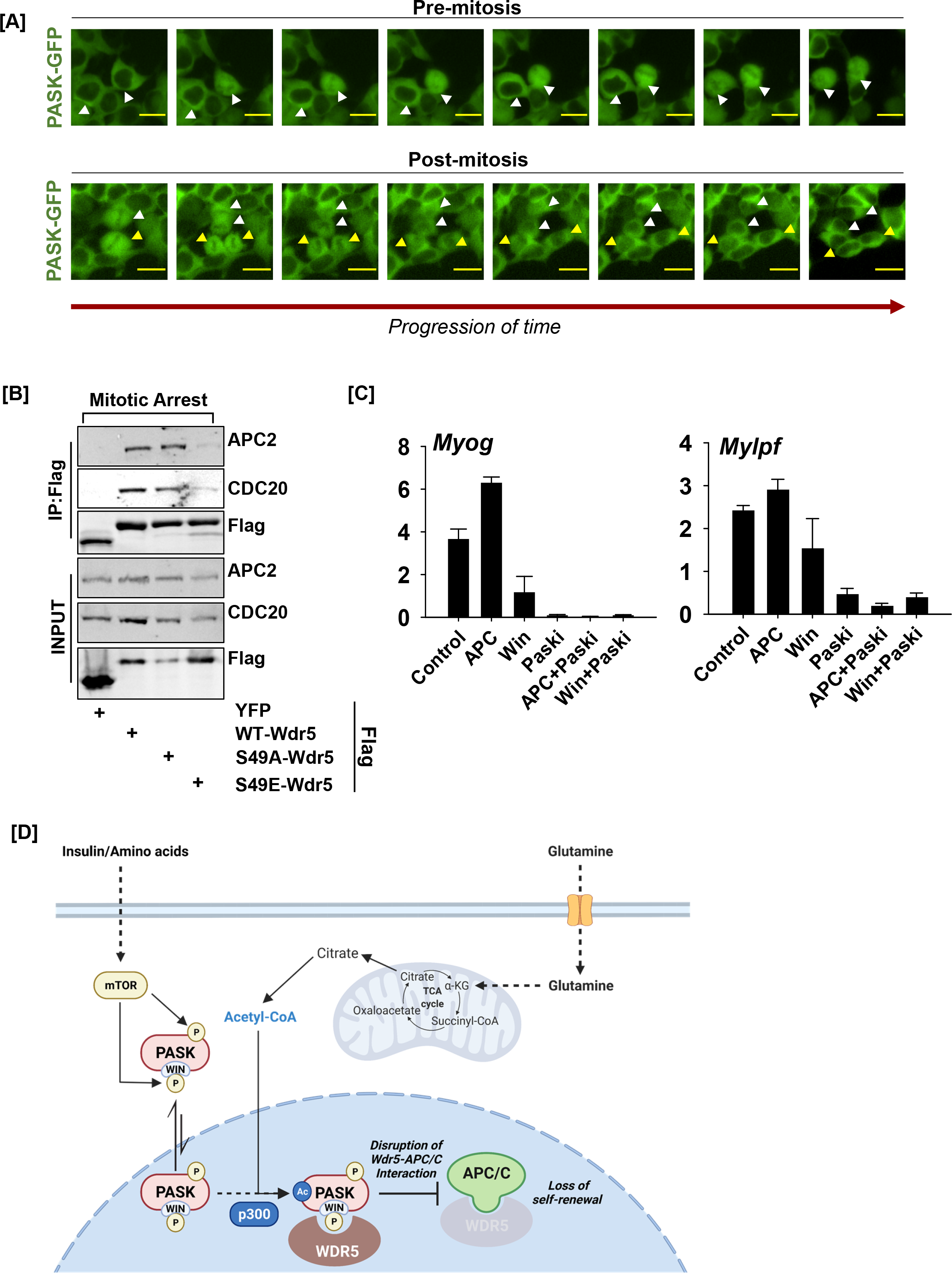
Negative regulation of the WDR5-APC/C interaction by PASK drives myogenesis. (A) Nucleo-cytoplasmic shuttling of GFP-hPASK during mitosis in asynchronous HEK293T cells showing possible nuclear entry of Pask in pre-mitotic cells prior to the nuclear membrane breakdown. (B) Phosphorylation of WDR5 at S49 disrupts the WDR5-APC/C interaction during mitosis. HEK-293T cells were transfected with FLAG-WDR5 (WT-WDR5), FLAG-WDR5 S49A (S49A-WDR5), FLAG-WDR5 S49E (S49E-WDR5), or YFP vector control (YFP), cultured in the presence or absence of glutamine as indicated, and treated with 50 μM PASK inhibitor (BioE-1197, PASKi). Cells were released from G2/M phase arrest into culture following nocodazole treatment. The lysate was immunoprecipitated with anti-FLAG beads prior to immunoblotting for APC2, CDC20, and FLAG. (C) Disruption of the WDR5-APC/C interaction by PASK is required for the expression of genes associated with skeletal muscle differentiation. Proliferating C2C12 myoblasts were treated with DMSO (control), proTAME (a selective inhibitor of APC/C, APCi), WIN inhibitor (WIN), PASKi (BioE-1197, PASKi), proTAME + PASKi (APC + PASKi), or WIN inhibitor + PASKi (WIN + PASKi). Transcript levels of *Myog* and *Mylpf* were quantified by RT-qPCR. (D) A model describing the mechanistic control of cell fate and identity by glutamine metabolism.

## References

1. Ahsan, S., Raval, M.H., Ederer, M., Tiwari, R., Chareunsouk, A., and Rodgers, J.T. (2020). Metabolism of glucose and glutamine is critical for skeletal muscle stem cell activation. bioRxiv, 2020.2007.2028.225847.

2. Andres, D.A., Young, L.E.A., Veeranki, S., Hawkinson, T.R., Levitan, B.M., He, D., Wang, C., Satin, J., and Sun, R.C. (2020). Improved workflow for mass spectrometry-based metabolomics analysis of the heart. J Biol Chem 295, 2676–2686.

3. Arnold, P.K., Jackson, B.T., Paras, K.I., Brunner, J.S., Hart, M.L., Newsom, O.J., Alibeckoff, S.P., Endress, J., Drill, E., Sullivan, L.B., and Finley, L.W.S. (2022). A non-canonical tricarboxylic acid cycle underlies cellular identity. Nature 603, 477–481.

4. Baksh, S.C., Todorova, P.K., Gur-Cohen, S., Hurwitz, B., Ge, Y., Novak, J.S.S., Tierney, M.T., Dela Cruz-Racelis, J., Fuchs, E., and Finley, L.W.S. (2020). Extracellular serine controls epidermal stem cell fate and tumour initiation. Nat Cell Biol 22, 779–790.

5. Banaszynski, L.A., Wen, D., Dewell, S., Whitcomb, S.J., Lin, M., Diaz, N., Elsasser, S.J., Chapgier, A., Goldberg, A.D., Canaani, E., et al. (2013). Hira-dependent histone H3.3 deposition facilitates PRC2 recruitment at developmental loci in ES cells. Cell 155, 107–120.

6. Chakrabarty, R.P., and Chandel, N.S. (2021). Mitochondria as Signaling Organelles Control Mammalian Stem Cell Fate. Cell Stem Cell 28, 394–408.

7. Chakraborty, A.A., Laukka, T., Myllykoski, M., Ringel, A.E., Booker, M.A., Tolstorukov, M.Y., Meng, Y.J., Meier, S.R., Jennings, R.B., Creech, A.L., et al. (2019). Histone demethylase KDM6A directly senses oxygen to control chromatin and cell fate. Science 363, 1217–1222.

8. Chang, J.S., Huypens, P., Zhang, Y., Black, C., Kralli, A., and Gettys, T.W. (2010). Regulation of NT- PGC-1alpha subcellular localization and function by protein kinase A-dependent modulation of nuclear export by CRM1. J Biol Chem 285, 18039–18050.

9. Choudhary, C., Weinert, B.T., Nishida, Y., Verdin, E., and Mann, M. (2014). The growing landscape of lysine acetylation links metabolism and cell signalling. Nature Reviews Molecular Cell Biology 15, 536–550.

10. Cutler, A.A., Pawlikowski, B., Wheeler, J.R., Betta, N.D., Elston, T., O’Rourke, R., Jones, K., and Olwin, B.B. (2021). The regenerating skeletal muscle niche guides muscle stem cell self-renewal. bioRxiv, 635805.

11. Danoviz, M.E., and Yablonka-Reuveni, Z. (2012). Skeletal muscle satellite cells: background and methods for isolation and analysis in a primary culture system. Methods Mol Biol 798, 21–52.

12. Das, S., Morvan, F., Morozzi, G., Jourde, B., Minetti, G.C., Kahle, P., Rivet, H., Brebbia, P., Toussaint, G., Glass, D.J., and Fornaro, M. (2017). ATP Citrate Lyase Regulates Myofiber Differentiation and Increases Regeneration by Altering Histone Acetylation. Cell Rep 21, 3003–3011.

13. Guarnaccia, A.D., Rose, K.L., Wang, J., Zhao, B., Popay, T.M., Wang, C.E., Guerrazzi, K., Hill, S., Woodley, C.M., Hansen, T.J., et al. (2021). Impact of WIN site inhibitor on the WDR5 interactome. Cell Rep 34, 108636.

14. He, S., Nakada, D., and Morrison, S.J. (2009). Mechanisms of stem cell self-renewal. Annu Rev Cell Dev Biol 25, 377–406.

15. Karakkat, J.V., Kaimala, S., Sreedharan, S.P., Jayaprakash, P., Adeghate, E.A., Ansari, S.A., Guccione, E., Mensah-Brown, E.P.K., and Starling Emerald, B. (2019). The metabolic sensor PASK is a histone 3 kinase that also regulates H3K4 methylation by associating with H3K4 MLL2 methyltransferase complex. Nucleic Acids Res 47, 10086-10103.

17. Kikani, C.K., Verona, E.V., Ryu, J., Shen, Y., Ye, Q., Zheng, L., Qian, Z., Sakaue, H., Nakamura, K., Du, J., et al. (2012). Proliferative and antiapoptotic signaling stimulated by nuclear-localized PDK1 results in oncogenesis. Sci Signal 5, ra80.

18. Kikani, C.K., Wu, X., Fogarty, S., Kang, S.A.W., Dephoure, N., Gygi, S.P., Sabatini, D.M., and Rutter, J. (2019). Activation of PASK by mTORC1 is required for the onset of the terminal differentiation program. Proc Natl Acad Sci U S A 116, 10382–10391.

19. Kikani, C.K., Wu, X., Paul, L., Sabic, H., Shen, Z., Shakya, A., Keefe, A., Villanueva, C., Kardon, G., Graves, B., et al. (2016). Pask integrates hormonal signaling with histone modification via Wdr5 phosphorylation to drive myogenesis. Elife 5.

20. Kırlı, K., Karaca, S., Dehne, H.J., Samwer, M., Pan, K.T., Lenz, C., Urlaub, H., and Görlich, D. (2015). A deep proteomics perspective on CRM1-mediated nuclear export and nucleocytoplasmic partitioning. eLife 4, e11466.

21. Lasko, L.M., Jakob, C.G., Edalji, R.P., Qiu, W., Montgomery, D., Digiammarino, E.L., Hansen, T.M., Risi, R.M., Frey, R., Manaves, V., et al. (2017). Discovery of a selective catalytic p300/CBP inhibitor that targets lineage-specific tumours. Nature 550, 128–132.

22. Lim, M.A., Kikani, C.K., Wick, M.J., and Dong, L.Q. (2003). Nuclear translocation of 3’- phosphoinositide-dependent protein kinase 1 (PDK-1): a potential regulatory mechanism for PDK-1 function. Proc Natl Acad Sci U S A 100, 14006–14011.

23. Liu, L., Cheung, T.H., Charville, G.W., Hurgo, B.M., Leavitt, T., Shih, J., Brunet, A., and Rando, T.A. (2013). Chromatin modifications as determinants of muscle stem cell quiescence and chronological aging. Cell Rep 4, 189–204.

24. Liu, L., Michowski, W., Kolodziejczyk, A., and Sicinski, P. (2019). The cell cycle in stem cell proliferation, pluripotency and differentiation. Nat Cell Biol 21, 1060–1067.

25. Liu, Y., Pelham-Webb, B., Di Giammartino, D.C., Li, J., Kim, D., Kita, K., Saiz, N., Garg, V., Doane, A., Giannakakou, P., et al. (2017). Widespread Mitotic Bookmarking by Histone Marks and Transcription Factors in Pluripotent Stem Cells. Cell Rep 19, 1283–1293.

26. Lu, V., Roy, I.J., and Teitell, M.A. (2021). Nutrients in the fate of pluripotent stem cells. Cell Metabolism 33, 2108–2121.

27. Mootha, V.K., Lindgren, C.M., Eriksson, K.-F., Subramanian, A., Sihag, S., Lehar, J., Puigserver, P., Carlsson, E., Ridderstråle, M., Laurila, E., et al. (2003). PGC-1α-responsive genes involved in oxidative phosphorylation are coordinately downregulated in human diabetes. Nature Genetics 34, 267–273.

28. Oh, E., Mark, K.G., Mocciaro, A., Watson, E.R., Prabu, J.R., Cha, D.D., Kampmann, M., Gamarra, N., Zhou, C.Y., and Rape, M. (2020). Gene expression and cell identity controlled by anaphase-promoting complex. Nature 579, 136–140.

29. Pauklin, S., and Vallier, L. (2013). The cell-cycle state of stem cells determines cell fate propensity. Cell 155, 135–147.

30. Pelham-Webb, B., Polyzos, A., Wojenski, L., Kloetgen, A., Li, J., Di Giammartino, D.C., Sakellaropoulos, T., Tsirigos, A., Core, L., and Apostolou, E. (2021). H3K27ac bookmarking promotes rapid post-mitotic activation of the pluripotent stem cell program without impacting 3D chromatin reorganization. Mol Cell 81, 1732–1748 e1738.

31. Puri, P.L., Avantaggiati, M.L., Balsano, C., Sang, N., Graessmann, A., Giordano, A., and Levrero, M. (1997a). p300 is required for MyoD-dependent cell cycle arrest and muscle-specific gene transcription. EMBO J 16, 369–383.

32. Puri, P.L., Sartorelli, V., Yang, X.J., Hamamori, Y., Ogryzko, V.V., Howard, B.H., Kedes, L., Wang, J.Y., Graessmann, A., Nakatani, Y., and Levrero, M. (1997b). Differential roles of p300 and PCAF acetyltransferases in muscle differentiation. Mol Cell 1, 35–45.

33. Sartorelli, V., Puri, P.L., Hamamori, Y., Ogryzko, V., Chung, G., Nakatani, Y., Wang, J.Y., and Kedes, L. (1999). Acetylation of MyoD directed by PCAF is necessary for the execution of the muscle program. Mol Cell 4, 725–734.

34. Shang, M., Cappellesso, F., Amorim, R., Serneels, J., Virga, F., Eelen, G., Carobbio, S., Rincon, M.Y., Maechler, P., De Bock, K., et al. (2020). Macrophage-derived glutamine boosts satellite cells and muscle regeneration. Nature 587, 626–631.

35. Subramanian, A., Tamayo, P., Mootha, V.K., Mukherjee, S., Ebert, B.L., Gillette, M.A., Paulovich, A., Pomeroy, S.L., Golub, T.R., Lander, E.S., and Mesirov, J.P. (2005). Gene set enrichment analysis: A knowledge-based approach for interpreting genome-wide expression profiles. Proceedings of the National Academy of Sciences 102, 15545–15550.

36. Sun, R.C., Dukhande, V.V., Zhou, Z., Young, L.E.A., Emanuelle, S., Brainson, C.F., and Gentry, M.S. (2019). Nuclear Glycogenolysis Modulates Histone Acetylation in Human Non-Small Cell Lung Cancers. Cell Metab 30, 903–916 e907.

37. TeSlaa, T., Chaikovsky, A.C., Lipchina, I., Escobar, S.L., Hochedlinger, K., Huang, J., Graeber, T.G., Braas, D., and Teitell, M.A. (2016). α-Ketoglutarate Accelerates the Initial Differentiation of Primed Human Pluripotent Stem Cells. Cell Metab 24, 485–493.

38. Wong, D.J., Liu, H., Ridky, T.W., Cassarino, D., Segal, E., and Chang, H.Y. (2008). Module Map of Stem Cell Genes Guides Creation of Epithelial Cancer Stem Cells. Cell Stem Cell 2, 333–344.

39. Wray, J., Kalkan, T., Gomez-Lopez, S., Eckardt, D., Cook, A., Kemler, R., and Smith, A. (2011). Inhibition of glycogen synthase kinase-3 alleviates Tcf3 repression of the pluripotency network and increases embryonic stem cell resistance to differentiation. Nat Cell Biol 13, 838–845.

40. Wu, X., Romero, D., Swiatek, W.I., Dorweiler, I., Kikani, C.K., Sabic, H., Zweifel, B.S., McKearn, J., Blitzer, J.T., Nickols, G.A., and Rutter, J. (2014). PAS kinase drives lipogenesis through SREBP-1 maturation. Cell Rep 8, 242–255.

41. Xu, D., Marquis, K., Pei, J., Fu, S.C., Cağatay, T., Grishin, N.V., and Chook, Y.M. (2015). LocNES: a computational tool for locating classical NESs in CRM1 cargo proteins. Bioinformatics 31, 1357–1365.

42. Xu, J., McPartlon, M., and Li, J. (2021). Improved protein structure prediction by deep learning irrespective of co-evolution information. Nature Machine Intelligence 3, 601–609.

43. Yang, C., Ko, B., Hensley, Christopher T., Jiang, L., Wasti, Ajla T., Kim, J., Sudderth, J., Calvaruso, Maria A., Lumata, L., Mitsche, M., et al. (2014). Glutamine Oxidation Maintains the TCA Cycle and Cell Survival during Impaired Mitochondrial Pyruvate Transport. Molecular Cell 56, 414–424.

44. Yu, Y., Newman, H., Shen, L., Sharma, D., Hu, G., Mirando, A.J., Zhang, H., Knudsen, E., Zhang, G.F., Hilton, M.J., and Karner, C.M. (2019). Glutamine Metabolism Regulates Proliferation and Lineage Allocation in Skeletal Stem Cells. Cell Metab 29, 966–978 e964.

